# Multiplexed single-cell profiling of post-perturbation transcriptional responses to define cancer vulnerabilities and therapeutic mechanism of action

**DOI:** 10.1101/868752

**Authors:** James M. McFarland, Brenton R. Paolella, Allison Warren, Kathryn Geiger-Schuller, Tsukasa Shibue, Michael Rothberg, Olena Kuksenko, Andrew Jones, Emily Chambers, Danielle Dionne, Samantha Bender, Brian M. Wolpin, Mahmoud Ghandi, Itay Tirosh, Orit Rozenblatt-Rosen, Jennifer A. Roth, Todd R. Golub, Aviv Regev, Andrew J. Aguirre, Francisca Vazquez, Aviad Tsherniak

**Affiliations:** Broad Institute of MIT and Harvard, Cambridge, MA, USA; Klarman Cell Observatory, Broad Institute of MIT and Harvard, Cambridge, MA, USA; Harvard Medical School, Boston, MA, USA; Brigham and Women’s Hospital, Boston, MA, USA; Department of Medical Oncology, Dana Farber Cancer Institute, Boston, MA, USA; Department of Molecular Cell Biology, Weizmann Institute of Science, Rehovot, Israel; Department of Pediatric Oncology, Dana Farber Cancer Institute, Boston, MA, USA; Howard Hughes Medical Institute, Chevy Chase, MD, USA; Koch Institute of Integrative Cancer Research, Cambridge, MA, USA; Department of Biology, MIT, Cambridge, MA, USA

## Abstract

Assays to study cancer cell responses to pharmacologic or genetic perturbations are typically restricted to using simple phenotypic readouts such as proliferation rate or the expression of a marker gene. Information-rich assays, such as gene-expression profiling, are generally not amenable to efficient profiling of a given perturbation across multiple cellular contexts. Here, we developed MIX-Seq, a method for multiplexed transcriptional profiling of post-perturbation responses across a mixture of samples with single-cell resolution, using SNP-based computational demultiplexing of single-cell RNA-sequencing data. We show that MIX-Seq can be used to profile responses to chemical or genetic perturbations across pools of 100 or more cancer cell lines, and combine it with Cell Hashing to further multiplex additional experimental conditions, such as multiple post-treatment time points or drug doses. Analyzing the high-content readout of scRNA-seq reveals both shared and context-specific transcriptional response components that can identify drug mechanism of action and can be used to predict long-term cell viability from short-term transcriptional responses to treatment.

## Introduction

Large-scale screens of chemical and genetic vulnerabilities across hundreds of cancer cell lines are important for identifying new therapeutic targets and are providing key insights into cancer biology and gene function^1–7^. However, the ability of these approaches to reveal the cellular mechanisms and pathways underlying such cancer vulnerabilities is typically limited by their reliance on a single readout of cell viability to assess the effects of each perturbation.

In contrast, information-rich, high-content readouts may provide opportunities to capture a more detailed picture of the cellular effects of a perturbation that underlie an observed fitness effect, or arise independently of any observable fitness effects^8–11^. In particular, expression profiles are a robust and informative measure for characterizing cellular responses to perturbations, with applications such as identifying drug mechanism of action (MoA), gene function, and gene regulatory networks^8–12^. High-throughput gene expression profiling in a limited number of contexts^10,13,14^ has been used to produce large datasets of perturbation signatures – most notably the Connectivity Map (CMAP)^10^ – enabling systematic analysis of the space of transcriptional responses across perturbations.

Until recently, however, such assays required each perturbation or cell type to be profiled separately, limiting the cost-effectiveness and broader adoption of gene expression profiling. In particular, previous efforts have largely focused on studying responses in a small number of cell line contexts. However, the response to perturbation is very often context specific, reflecting the interaction between the perturbation and a cell’s particular genomic or functional features. For example, targeted drugs may elicit responses only in cell lines harboring particular oncogenic mutations, or expressing certain genes, making observed results specific to the particular cell line models chosen^4,6,7,15,16^. More generally, the inability to efficiently measure transcriptional responses across diverse cell contexts has limited our understanding of how perturbation effects differ across the broad range of genomic and molecular cell states, which could be critical for predicting the therapeutic response of patient tumors. The recent advent of single-cell genomics^17,18^, and development of methods for profiling cell viability in pooled cell cultures^19^ could together help address these challenges. In parallel, new assays, such as Perturb-Seq^8,9^, have combined pooled perturbation screens with a single-cell RNA-Seq (scRNA-seq) readout and could thus provide the necessary scale and resolution to assay many cells within a mixed culture.

To facilitate the study of post-treatment gene expression signatures across multiple cell lines in parallel, we have developed **MIX-Seq**: **<u>M</u>**ultiplexed **<u>I</u>**nterrogation of gene e**<u>X</u>**pression through single-cell RNA **<u>Seq</u>**uencing. This approach leverages the ability to pool hundreds of cancer cell lines and co-treat them with one or more perturbations. We then apply scRNA-seq to simultaneously profile the cells’ responses and resolve the identity of each cell based on single nucleotide polymorphism (SNP) profiles. MIX-Seq enables efficient study of transcriptional signatures after pharmacologic or genetic perturbation, evaluation of temporal evolution of post-perturbation transcriptional response, investigation of the mechanism-of-action (MoA) of novel small-molecule compounds and development of novel therapeutic response prediction methods in cancer cell models.

## Results

### MIX-Seq: Multiplexed cell line transcriptional profiling using scRNA-seq

MIX-Seq uses scRNA-seq to measure the transcriptional effects of a perturbation across diverse cancer cell lines grown and perturbed in a pool (**Fig. 1a**). Specifically, we co-culture cancer cell lines in pools and treat them with a small molecule compound (or genetic perturbation), following our PRISM approach^19^. To ascertain transcriptional response signatures, cell-specific transcriptomes are measured using scRNA-seq after a defined time interval following perturbation. To assign each profiled cell to its respective cell line, we created a computational demultiplexing method that classifies cells by their genetic fingerprints. Specifically, we first estimated allelic fractions across a panel of commonly occurring SNPs for a set of reference cell lines (**Supplementary Table 1**). For each single cell, we then determine the reference cell line with the highest likelihood of generating the observed pattern of SNP reads, using a generalized linear model (see Methods). This approach also allows for identification of multiplets of co-encapsulated cells^20^, where two or more cells from different cell lines are unintentionally tagged with the same cell barcode during droplet-based single-cell library preparation, and it provides quality metrics that can be used to identify and remove low-quality cells (**Supplementary Fig. 1**), such as empty droplets^18,21^.

**Fig. 1:**
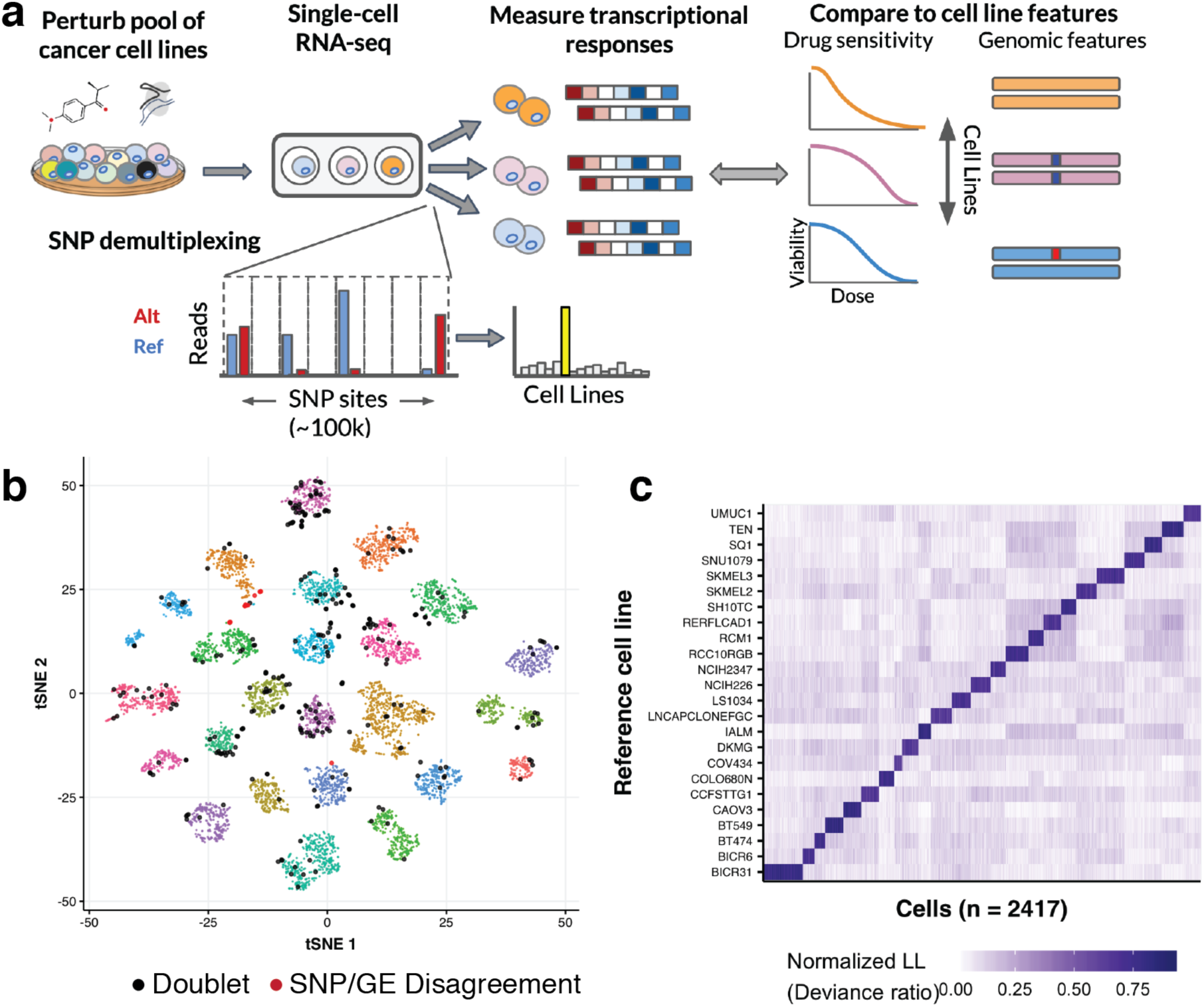
MIX-Seq platform leverages SNP-based demultiplexing of mixtures of single-cell transcriptional profiles. **a)** Schematic diagram illustrating the MIX-Seq platform. **b)** t-SNE representation of expression profiles from an example negative-control experiment (combination of DMSO-treated and untreated cells), showing clear clustering by cell line (color indicates SNP-based parental cell line classification). Black dots show cells classified as doublets. Red dots indicate cells where gene expression and SNP-based classifications disagree (0.2% of singlet cells, 16/6926). **c)** Heatmap showing likelihoods assigned by the SNP-classification model for each cell coming from each parental cell line. The model consistently picks out a single cell line, from among the 24 ‘in-pool’ cell lines, with high confidence.

We confirmed the classification accuracy of our SNP-based demultiplexing approach on scRNA-seq profiles from pools of either 24 or 99 cancer cell lines at baseline (either untreated, or treated with a vehicle control DMSO) in two ways. First, we classified cell identities independently by either their gene expression profiles (see Methods) or their SNPs. After removal of low-quality cells and doublets (**Supplementary Fig. 1**), we found that classifications based on SNP and gene expression profiles were very similar (> 99% agreement; **Fig. 1b; Supplementary Fig. 2**). While either feature could be thus used to accurately classify cell identities; we focus on SNP-based classification here as it is inherently robust to perturbations that could dramatically alter the cells’ gene expression profiles, and could be applied to pools of primary cells of the same type from different individuals (e.g. iPS cells).

Second, we estimated the error rate of SNP-based classification by assessing the frequency with which the model selected cell lines that were not in the experimental pools. Out of a reference panel of 494 cell lines, the model consistently selected a cell line from within the experimental pool with high confidence (**Fig. 1c**), and never picked an out-of-pool cell line (0/84,869 cells were classified ‘out of pool’). Notably, though we tested MIX-Seq with experimental pools of up to 99 cell lines, these analyses show that SNP profiles can be used to distinguish among much larger (>500) cell line pools. Furthermore, based on downsampling analysis, we found that SNP-based cell classifications can be applied robustly to cells with as few as 50-100 detected SNP sites^20^ (**Supplementary Fig. 3**). Our SNP-based model also detected doublets at the expected rate (based on the cell loading densities)^22^, and consistently identified both cells of doublet pairs among those in the experimental pool (**Supplementary Fig. 1**).

In summary, these results show that MIX-Seq allows for multiplexed transcriptional profiling across pools of cancer cell lines using scRNA-seq, with robust SNP-based demultiplexing and doublet detection that can be scaled to large pools of cell lines.

### MIX-seq identifies selective perturbation responses and MOA

Next, we evaluated whether MIX-Seq could distinguish biologically meaningful changes in gene expression in the context of drug treatment, including identifying differential cell line responses related to an established biomarker, and recovering the compounds’ MoA from the measured transcriptional responses. We treated pools of well-characterized cancer cell lines with 13 drugs, followed by scRNA-seq at 6 and/or 24 hours after treatment (**Supplementary Table 2**). These included 8 targeted cancer therapies with known mechanisms, 4 pan-lethal compounds that broadly kill most cell lines, and one tool compound (BRD-3379) with unknown MoA, which was found to induce strong selective killing in a high-throughput screen. We compared our scRNA-seq based phenotyping to long-term viability responses measured for these drugs and cell lines from the genomics of drug sensitivity in cancer (GDSC) screening dataset^4,7^, as well as data generated using the PRISM assay^16,19^ (see Methods).

As an example, we first consider nutlin, a selective MDM2 inhibitor, which we applied to a pool of 24 cell lines. MDM2 is a negative regulator of the tumor suppressor gene TP53, and nutlin is known to elicit rapid apoptosis and cell cycle arrest exclusively in cell lines that have wild-type (WT) TP53^23^. Jointly embedding the expression profiles of 7,317 single cells treated with either nutlin or vehicle control (DMSO) in 2D-space revealed clear clustering by cell line, with robust shifts in the nutlin-treated cell populations for some cell lines, but not others (**Fig. 2a**). Estimating the average drug-induced change in gene abundance for each cell line (see Methods) revealed a robust response in each of the 7 TP53 WT cell lines in the pool, but only minimal changes in cell lines harboring TP53 mutations, as expected (**Fig. 2b-d**). Furthermore, pathway enrichment analysis (see Methods) of the average transcriptional response among TP53 WT cell lines showed clear up-regulation of genes in the TP53 downstream pathway, as well as down-regulation of cell cycle processes (**Fig. 2e**). Thus, the differential transcriptional programs identified by MIX-Seq can be used to inform a compound’s MoA.

**Fig. 2:**
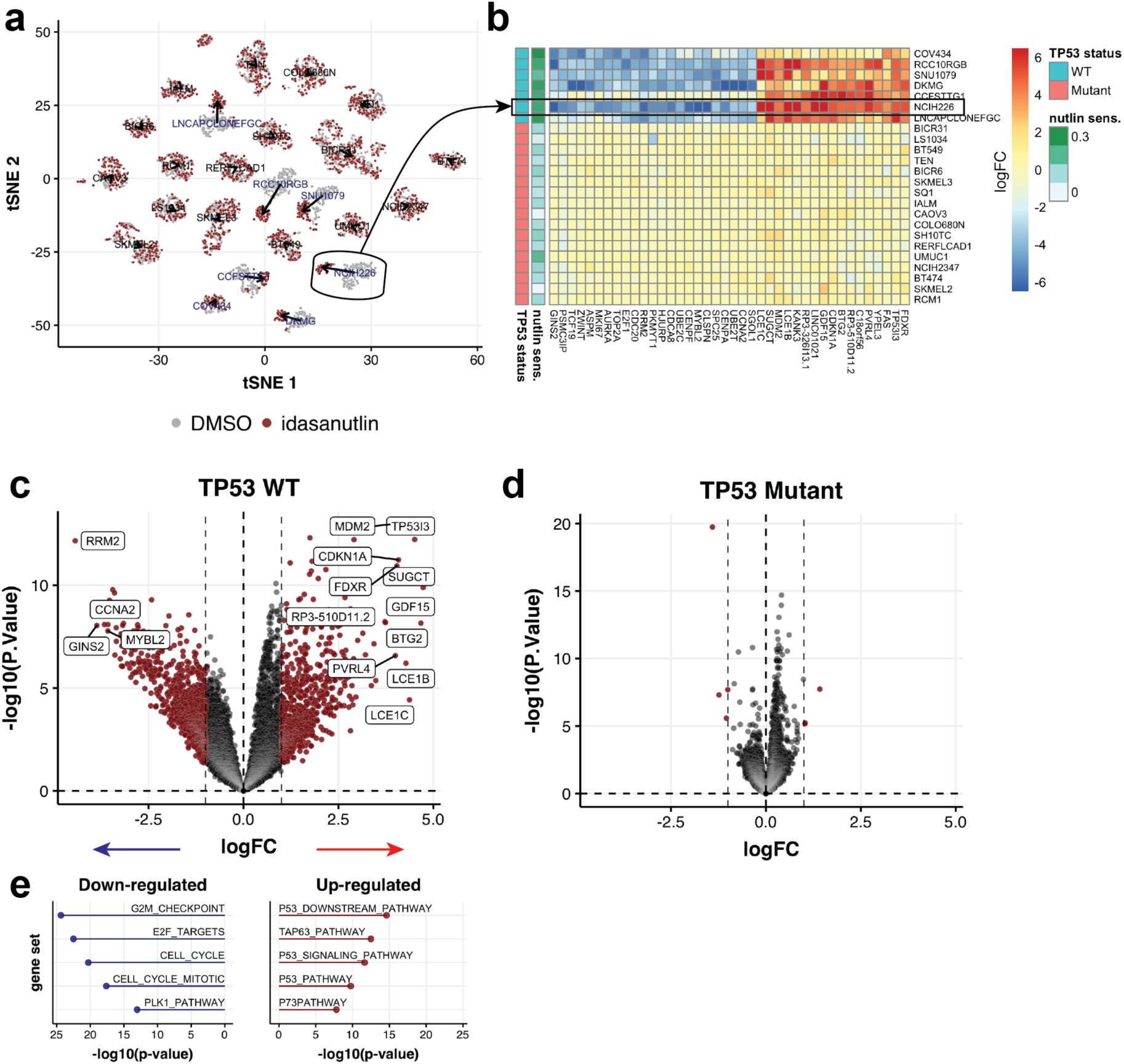
Identification of selective transcriptional responses and compound MoA. **a)** t-SNE representation of cells treated with DMSO control (blue) or nutlin (red) across a pool of 24 cell lines. Arrows indicate the shift in the population median coordinates for each cell line. **b)** Heatmap showing average log fold-change estimates for each cell line for top differentially expressed genes. Nutlin sensitivity is given by 1 – area under dose response curve (AUC, see Methods). **c)** Volcano plot showing strong gene expression changes in response to nutlin treatment across TP53 WT cell lines (n = 7). p-values were estimated based on empirical-Bayes moderated t-statistics, using the limma-trend pipeline^29,30^. **d)** Same as **c** for TP53 mutant cell lines (n = 17), showing little gene expression change in response to nutlin treatment. **e)** Gene set analysis identifies gene sets that are up-(right) and down-(left) regulated by nutlin treatment in the TP53 WT cell lines.

Across nearly all 13 drugs profiled, we were able to identify robust transcriptional response signatures, and these signatures were often highly informative about the compounds’ MoA^24,25^. For example, treatment with the proteasome inhibitor bortezomib elicited strong up-regulation of protein folding and heat-shock response pathways (**Supplementary Fig. 4a**). Gemcitabine, a chemotherapy drug, altered expression of apoptosis-related genes (**Supplementary Fig. 4b**), and mTOR signaling was the top down-regulated gene set following treatment with the mTOR inhibitor everolimus (**Supplementary Fig. 4c**). For the two inhibitors of anti-apoptotic proteins (navitoclax, and the MCL1i AZD-5591) we observed relatively weak transcriptional responses without clear cell line selectivity, as discussed further below. Taken together, these results demonstrate the ability of MIX-Seq to measure selective transcriptional effects of a drug across a pool of cell lines, and highlight the utility of such information for identifying a drug’s cellular effects and MoA^10,12^.

In addition to measuring drug-responses, MIX-Seq can also be used to study the transcriptional effects of genetic perturbations in a pool of cell lines. For example, we introduced two sgRNAs targeting the gene glutathione peroxidase 4 (GPX4) by lentiviral transduction into a pool of 50 cell lines, and performed scRNA-seq at either 72 or 96 hours post-infection (see Methods). GPX4, a gene involved in lipid metabolism, was selected in part because it represents a strong selective dependency among cell lines with a mesenchymal phenotype^26^, and is highly and ubiquitously expressed, allowing us to directly assess the level of on-target knockdown of GPX4 mRNA^8^. Both GPX-targeting guides produced robust on-target reduction of GPX4 mRNA compared to control guides (**Supplementary Fig. 5a**) across each of the 49 cell lines detected in the pool (**Supplementary Fig. 5b,c**). The up-regulated genes included EEF1A2, shown to play a role in regulating lipid metabolism^27^, and genes involved in glycan-synthesis, which has been reported to regulate glutathione levels^28^ (**Supplementary Fig. 5d**). Notably, however, we did not observe clear differences in the transcriptional responses of cell lines known to be dependent vs. non-dependent on GPX4 CRISPR viability screens^2^ (**Supplementary Fig. 5e**).

### Single-cell profiling enables characterization of heterogeneous population responses

The single-cell resolution from MIX-Seq further allows us to determine how perturbations affect the composition and state of heterogeneous cell populations. For instance, we can infer the cell cycle phase of each cell from its expression profile^31^, and then determine how perturbations impact the cell cycle phase composition of different cell lines. Applying this approach showed that nutlin treatment elicited a pronounced G0/G1-arrest phenotype selectively among the TP53 WT cell lines (**Fig. 3a,b**), as expected.

**Fig. 3:**
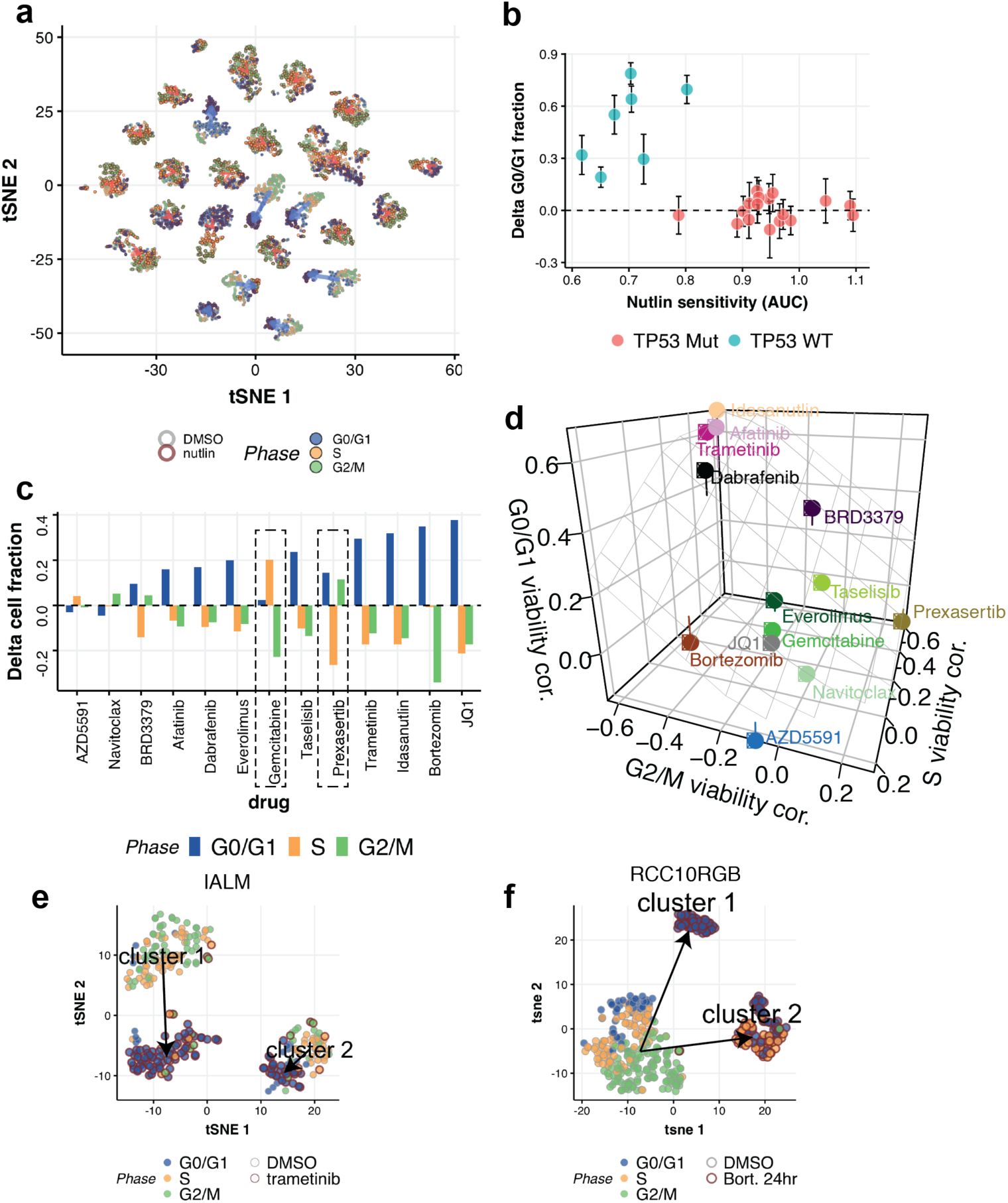
Single-cell resolution enables identification of heterogeneity in pre- and post-perturbation transcriptional programs. **a)** t-SNE representation of gene expression profiles of DMSO (light gray outline) and nutlin-treated (dark red outline) cell populations for a pool of 24 cell lines (as in **Fig. 2**). Cells are colored by their inferred cell cycle phase. TP53 WT cell lines (blue arrows) show predominance of G0/G1-phase cells after nutlin treatment not observed in TP53 mutant cell lines (red arrows). **b)** Quantification of the change in proportion of cells in G0/G1 shows that nutlin treatment elicits G1-arrest selectively among the TP53 WT cell lines. Error bars show the 95% CI. **c)** Average change in the fraction of cells in each cell cycle phase for each drug treatment (averages are weighted by measured drug sensitivity, all for 24 hr time points). **d)** Comparison of the Pearson correlation between measured drug sensitivity and the changes in G2/M-, G0/G1-, and S-phase cell proportions for all compounds. Lattice shows the regression plane of the z-coordinate **e)** 2D representation of IALM single-cell expression profiles after DMSO (gray stroke) or trametinib (blue stroke) treatment (24 hours). Inferred cell cycle phase depicted by fill color. **f)** Similar to **e**, showing two sub-populations of RCC10RGB cells emerging 24 hours after treatment with bortezomib

We next systematically assessed the average effects of each compound on cell cycle phase composition (**Fig. 3c, Methods**). At 24 hours post-treatment, most drugs produced an increase in the proportion of cells in G0/G1 (10/13 drugs) and concomitantly decreased the proportion of cells in S (9/13) and G2/M phases (9/13), consistent with cell cycle arrest at the G1/S transition (**Fig. 3c**). Two notable exceptions were the DNA-damaging agent gemcitabine and the CHEK1/2 inhibitor prexasertib. Gemcitabine also decreased the proportion of cells in G2/M but with an increase in S-phase cells, consistent with its known role in triggering CHEK1-mediated S-phase arrest. Prexasertib decreased the proportion of S-phase cells, and slightly increased the fraction of G2/M cells, consistent with inhibition of CHEK-1 mediated DNA-damage checkpoints leading to dis-regulated progression of cells through the cell cycle^32^.

For compounds that are selectively active in only some cell lines, cell cycle effects were well-correlated with their measured viability effects across the cell lines, such that drugs typically had larger cell cycle effects in cell lines that were more sensitive (**Fig. 3d**). We also used the single-cell data to directly estimate drug-induced changes in relative cell abundance, finding that selective compounds consistently decreased the representation of more sensitive cell lines in the pool, particularly when measured 24 hours post-treatment (**Supplementary Fig. 6**). Thus, MIX-Seq can reliably read out the effects of perturbations on cell cycle progression as well as overall cell viability.

We next leveraged the single-cell nature of MIX-Seq data to study how perturbations affect different sub-populations of cells, and potentially alter patterns of transcriptional heterogeneity within a cell line. Cancer cell lines exhibit substantial genetic^33–35^, epigenetic^31,36,37^, and transcriptional^35,36,38,39^ heterogeneity. For example, we found that the lung cancer cell line IALM was composed of two distinct subpopulations at baseline, and these subpopulations showed significant differences in their transcriptional response to trametinib (**Fig. 3e; Supplementary Fig. 7a,b**). In another example, the kidney cancer cell line RCC10RGB consisted of a single population at baseline, but treatment with bortezomib gave rise to two distinct cell populations that were distinguished by expression of cell cycle and DNA damage response genes (**Fig. 3f; Supplementary Fig. 7c-e**), a pattern that was also observed in several other cell lines (**Supplementary Fig. 7f-i**).

These examples highlight the ability of MIX-Seq to not only identify differences in average perturbation responses across cell lines, but also to reveal the detailed effects of such perturbations on heterogeneous cell populations with single-cell resolution.

### Large-scale profiling identifies shared viability-related response signatures across drugs

Next, we wanted to understand the relationship between short-term transcriptional responses and long-term viability effects. While we initially profiled cells in a pool of 24 cell lines; we now leveraged SNP-demultiplexing in two much larger cell line pools of nearly 100 cell lines each (97 and 99) treated with 8 and 4 small molecules respectively. This allowed us to identify context-dependent response signatures, and short term differential responses related to long-term drug sensitivity (**Fig. 4a,b**). We employed a statistical modeling approach relating the (average) transcriptional changes measured in each cell line to their viability response in the drug sensitivity data from GDSC and PRISM^4,16,19^. Specifically, we decompose the change in expression of each gene into two components: a *viability-independent* response component (*β*_*0*_) characterizing the response of completely insensitive cell lines, and a *viability-related* response component (*β*_*1*_) characterizing the difference between sensitive and insensitive cell lines (**Fig. 4c**, top, Methods).

**Fig. 4:**
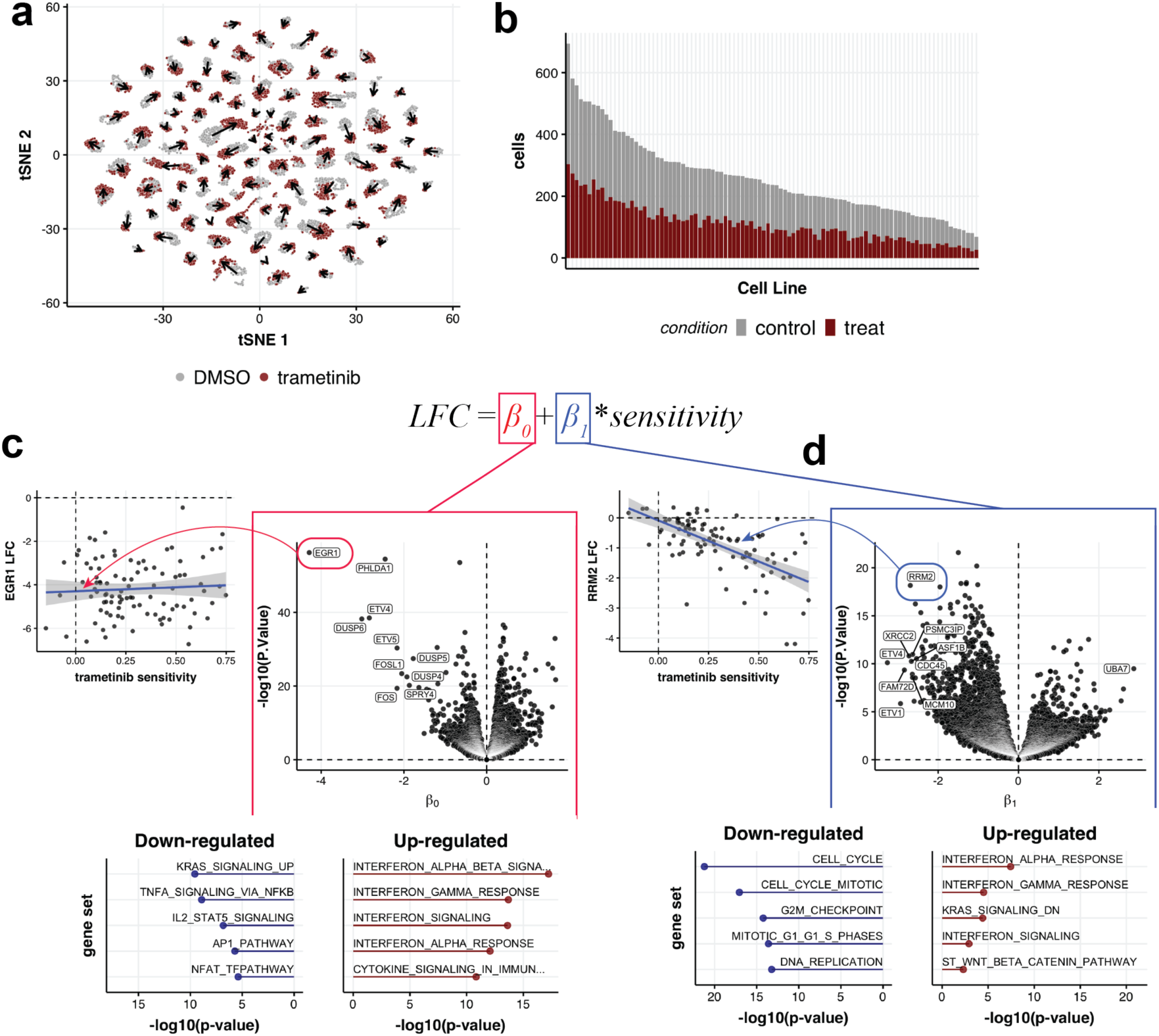
Post-perturbation transcriptional response signatures are comprised of distinct viability-related and -independent components. **a)** t-SNE representation of single-cell expression profiles in a 99 cell line pool treated with vehicle control (gray) or trametinib (red). Arrows indicate trametinib-induced shift of population median coordinates for each cell line. **b)** Histogram showing the number of cells recovered in each cell line and condition. **c)** Volcano plot showing the viability-independent response for each gene, representing the ‘y-intercept’ of a linear fit of expression change to drug sensitivity. Inset at left shows this relationship for an example gene: EGR1. Inset below shows top up- (right, red) and down-regulated (left, blue) gene sets. **d)** Same as **c** for the viability-related response

As an example, we first consider treatment of the 99 cell line pool with the MEK inhibitor trametinib, along with vehicle control (DMSO). We recovered more than 100 cells per cell line on average in each condition, detecting 97/99 cell lines with a minimum of 20 cells in each condition (average 130 cells/condition; **Fig. 4a,b**). Downsampling analysis suggested that measuring tens of cells per condition was sufficient to estimate each cell line’s transcriptional response profile (**Supplementary Fig. 8**).

The *viability-independent* response to trametinib included a strong down-regulation of MAPK signaling genes, including EGR1, ETV4/5, DUSP4/5/6, and SPRY2/4, the KRAS signaling pathways, and TNF-alpha signaling, as well as up-regulation of the interferon response (**Fig. 4c**), consistent with previous reports^40,41^. In contrast, the *viability-related* component showed strong down-regulation of cell-cycle processes (**Fig. 4d**). This result, along with the inferred G0/G1-arrest in sensitive cell lines (**Fig. 3c,d**), implicates a selective cell cycle arrest as mediating the long-term viability effects of trametinib.

Applying this analysis across all 8 selective compounds profiled with MIX-Seq, we found several core components of the viability-related response that were largely shared across compounds. These were highly enriched for cell-cycle genes, which were selectively down-regulated in the sensitive cell lines in virtually all the selective compounds profiled (**Supplementary Fig. 9**). A smaller set of genes related to translation were consistently up-regulated in sensitive cell lines, potentially owing to elevated translation in G1-arrested cells^42^. Notably, the shared signature was also apparent in cells treated with pan-toxic compounds, such as prexasertib, the BRD2-inhibitor JQ-1, bortezomib, and gemcitabine (**Supplementary Fig. 9**). This suggests that this is a general transcriptional signature of decreased cell viability and/or proliferation. Other viability-related response components were compound-specific, most notably a set of TP53-signaling related genes that were up-regulated in sensitive cell lines only with nutlin treatment (**Supplementary Fig. 9**). Finally, the two inhibitors of anti-apoptotic proteins -- navitoclax and AZD5591 -- were unique among the compounds tested in that they did not produce robust transcriptional response signatures despite having strong selective viability effects. Despite the lack of strong transcriptional response, AZD5591 (but not navitoclax) produced a clear depletion of cell abundance in the more sensitive cell lines at this time point (24 hr), and the magnitude of this cell line depletion was well-correlated with external measurements of drug-sensitivity (**Supplementary Fig. 6**).

In order to determine how the number of different cell lines profiled impacts estimation of these transcriptional response components we performed a downsampling analysis (**Supplementary Fig. 10**). While the average response across cell lines could be estimated reliably from relatively few (5-10) lines, estimates of the viability-related and viability-independent response components became more robust (as measured by their similarity to estimates using all cell lines) when including data from many lines (i.e. ∼50 or more) (**Supplementary Fig. 10**).

### Dual multiplexing of cell lines and time points highlights early onset of viability-independent programs and later onset of viability-related programs

Many perturbations will elicit cellular responses that evolve in complex ways over time, suggesting that more information could be obtained by profiling cells across an entire sequence of time points post-treatment. Several methods have recently been developed for introducing sample-specific barcodes to allow multiplexing of scRNA-seq measurements across experimental conditions^21,43^, such as multiple time points^43^. In particular, Cell Hashing^21^ uses oligonucleotide-conjugated antibodies against cell-surface antigens (called hashtags) to label cells with unique barcodes for each experimental condition. Since MIX-Seq uses naturally occurring “SNP barcodes” to multiplex cell lines, it can easily be combined with such approaches to allow for dual-multiplexing of cell lines and experimental conditions with a single scRNA-seq readout.

Leveraging this approach, we measured responses of a pool of 24 cell lines to trametinib along 5 time points, ranging from 3 to 48 hours post-treatment, using Cell Hashing to multiplex treatment conditions (**Fig. 5a**). As controls, we included DMSO-treated samples at each of the 5 time points, in addition to untreated samples, for a total of 11 conditions. Hashtag reads provided robust labeling of treatment conditions, with good tagging efficiency across all cell lines (**Supplementary Fig. 11**). Since we did not observe substantial differences in DMSO-treated cells across time points (**Supplementary Fig. 12**), we pooled them together for subsequent analysis, yielding a total of 13,713 clearly tagged single cells across all treatment conditions and cell lines.

**Fig. 5:**
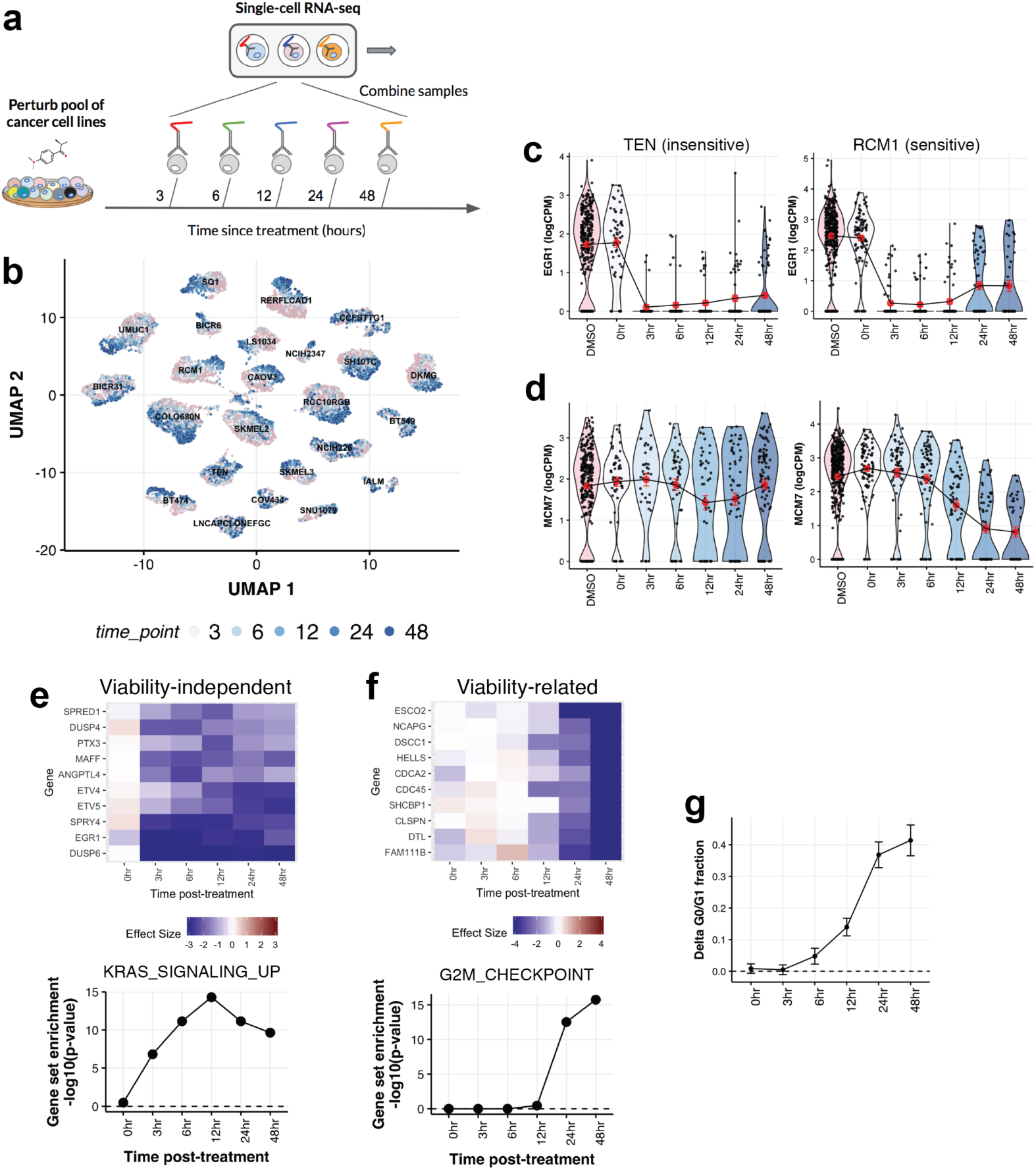
Temporal resolution of post-treatment transcriptional response using dual-multiplexing across cell lines and time points. **a)** Schematic diagram illustrating experiment using Cell Hashing to multiplex scRNA-seq of cell line pools sampled at different time points following drug treatment. **b)** UMAP plot showing 13,713 cells across a pool of 24 cell lines at different times following treatment with trametinib (shades of blue), or DMSO control (pink). **c)** Single-cell expression levels of EGR1 at different time points following trametinib treatment for an example insensitive/sensitive cell line (left/right). Red dots depict the mean expression levels at each time point, and error bars show the interval +/- s.e.m. **d)** Same as **c**, for MCM7. **e)** (Top) Time course of the viability-independent response for top down-regulated genes. (Bottom) Enrichment of HALLMARK_KRAS_SIGNALING_UP genes in the down-regulated viability-independent response at each time point. **f)** (Top) Same as **e**, showing time course of the viability-related response for top down-regulated genes. (Bottom) Enrichment of HALLMARK_G2M_CHECKPOINT genes in the viability-related response at each time post-treatment. **g)** Average time course of G0/G1-arrest across cell lines (n = 24). Error bars indicate interval +/- s.e.m.

The single-cell expression profiles illustrated strong time-dependent changes in response to trametinib treatment, whose magnitude varied considerably across cell lines (**Fig. 5b**). To better understand these changes, we examined the time courses of transcriptional changes for key trametinib-response genes. For example, EGR1, an immediate early response gene known to be activated by MAPK signaling^44^, was dramatically down-regulated 3 hours after trametinib treatment in both the sensitive cell line RCM1 and the insensitive line TEN (**Fig. 5c**). In contrast, MCM7, a cell-cycle-related gene that was part of the viability-related response, was selectively down-regulated only in the sensitive line RCM1, and only after 12-24 hours post-treatment (**Fig. 5d**).

We next applied our statistical model (**Fig. 4c,d**) to quantify the temporal evolution of *viability-related* and *viability-independent* components of the trametinib response for each gene, integrating across all cell lines. Down-regulated genes in the *viability-independent* response showed a range of temporal patterns, with several (such as EGR1 and DUSP6) reaching maximal down-regulation 3 hours post-treatment (**Fig. 5e**). In contrast, the *viability-related* response emerged much later, with genes such as GINS2, E2F1, and MCM4 showing selective down-regulation in sensitive cell lines only 12-24 hours post-treatment (**Fig. 5f**). The viability-independent down-regulation of the KRAS signaling pathway emerged 3 hours after treatment (**Fig. 5e**), while the viability-related down-regulation of cell cycle genes started 24 hours after treatment (**Fig. 5f**). The latter was also consistent with the time course of G0/G1-arrest based on inferred cell cycle phases (**Fig. 5g**).

These results thus highlight the utility of large-scale transcriptional profiling, both across cell lines and time points, to identify the different components of drug response. The ability to separate these transcriptional components could provide clues to both the initial effects of target-engagement, as well as the mechanism underlying selective loss of cell viability, and more powerfully inform MoA.

### Prediction of long-term viability responses from short-term post-treatment MIX-Seq profiles

Transcriptional profiling of drug responses across large panels of cell lines also enables the application of machine learning methods to discover different response patterns across cell lines without *a priori* knowledge of the relevant genomic/molecular features driving such differences. As a simple illustration of this, we first applied principal component analysis (PCA) to the matrix of trametinib responses across cell lines, measured 24 hours post-treatment (**Fig. 6a**). The first principal component (PC1), captured differences in trametinib sensitivity across cell lines (**Fig. 6b,c**). Indeed, across 9/13 tested drugs, PC1 of the transcriptional response matrix (measured at 24 hours post-treatment) was significantly correlated with the cell lines’ measured drug sensitivity (FDR < 0.1; **Supplementary Fig. 13**), suggesting that this is often a predominant source of response heterogeneity. For trametinib treatment, PC2 identified a pattern of differential response among a subset of trametinib-sensitive cell lines with high baseline expression of SOX10 (mostly BRAF mutant melanomas) (**Fig. 6d,e**). This example thus highlights the power of transcriptional profiling across cell contexts to identify multiple biologically-relevant factors underlying the differential cellular response to the drug.

**Fig. 6:**
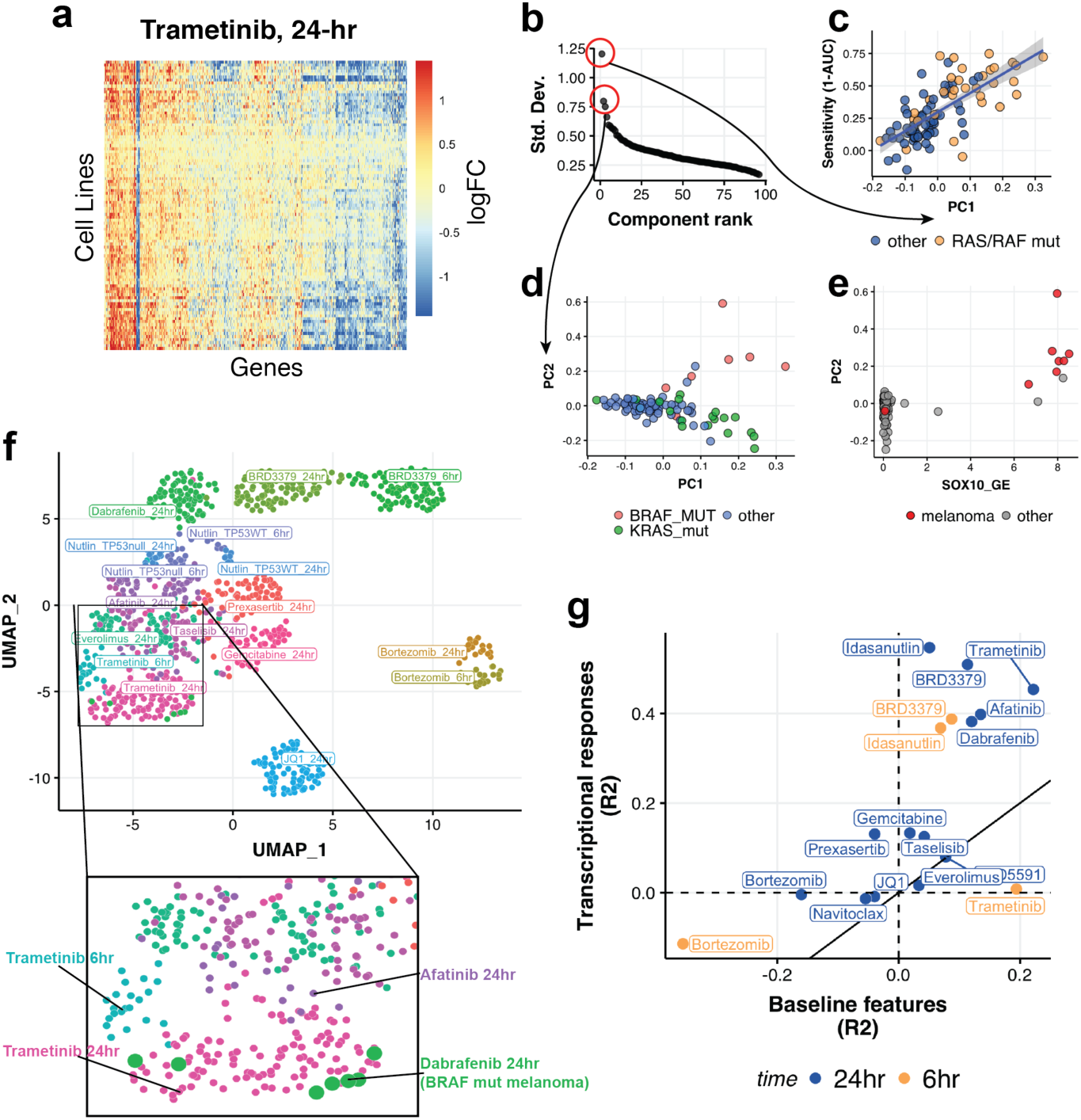
Machine learning analysis powered by large-scale transcriptional profiling. **a)** Matrix of measured transcriptional responses across cell lines 24 hr after trametinib treatment. **b)** Eigenvalue spectrum of PCA applied to the matrix from **a. c)** The projection of each cell line’s response onto PC1 is plotted against its measured trametinib sensitivity (yellow points indicate cell lines with activating RAS/RAF mutations). Linear regression trend line (with 95% CI interval) are shown in blue. **d)** Scatterplot of PC1 vs PC2 loadings across cell lines. While PC1 captures differences in trametinib sensitivity, PC2 largely captures the difference between KRAS mutant lines (green) and BRAF mutant melanomas (red). **e)** Comparison of PC2 scores with expression of the melonoma-specific transcription factor SOX10 across cell lines. **f)** UMAP embedding of transcriptional response profiles across drugs, cell lines, and post-treatment time points. Points are colored by treatment condition (drug and time point). Inset below shows zoomed view of the region indicated by the rectangle, with the larger green dots representing responses of BRAF mutant melanoma lines to the BRAFi dabrafenib. **g)** Accuracy of models trained to predict drug sensitivity using either measured transcriptional responses or baseline ‘omics’ features of the cell lines (using the same set of cell lines). Predictions based on transcriptional profiling at 6 and 24 hr post-treatment are indicated by the gold and blue dots respectively.

In order to identify, in an unsupervised manner, global patterns of transcriptional responses across cell lines, compounds and time points, we used UMAP^45^ to create a 2D map of all the combined perturbation response profiles. While perturbation response profiles mostly grouped by perturbation type (drug and post-treatment time point) (**Fig. 6f**), relationships between the set of responses for related perturbation types were also apparent. For example, responses to the same drug profiled at multiple post-treatment time points were nearby in UMAP space, and functionally related drugs such as taselisib (PIK3CAi) and everolimus (MTORi), as well as trametinib (MEKi) and afatinib (EGFRi), clustered near one another. Interestingly, the response of BRAF mutant cell lines to the BRAFi dabrafenib grouped with trametinib response profiles, rather than with the other dabrafenib responses (**Fig. 6f**).

Finally, we tested the feasibility of using short-term transcriptional responses to predict the long-term viability effects of a drug, which could have clinical applications in therapeutic response prediction, as patient cells can be transcriptionally profiled without long term cultures. We trained random forest models to predict long-term viability responses using the measured transcriptional response profiles for each cell line (across-cell averages). We evaluated the accuracy of the models using the R^2^ of predictions on held-out test cell lines (by 10-fold cross-validation; see Methods). For comparison, we also trained models to predict the viability response data using the baseline ‘omics’ features of the cell lines, including their baseline expression levels (from bulk RNA-seq data) and the presence of damaging or hotspot mutations^6,46^. Across drugs, transcriptional response signatures were more predictive of long-term viability responses compared to the cell lines’ baseline features (**Fig. 6g**; *n* = 17; *p* = 8.4×10^−4^; Wilcoxon signed rank test). This difference was particularly pronounced for the transcriptional responses measured 24 hours post-treatment, though we also obtained good predictive performance using 6-hour responses for some drugs. Notably, even when we used all available data to train models on the baseline features, such that they had access to much larger training samples (*e*.*g*., n = 741 vs 24 cell lines for nutlin), transcriptional response profiles still compared favorably for predicting viability effects for most drugs (**Supplementary Fig. 14**).

These results suggest that post-treatment transcriptional signatures can provide a robust signal of cellular response to drugs that could be applied to predict their long-term viability effects.

## Discussion

Here we present an experimental and computational platform (MIX-Seq) for performing highly multiplexed transcriptional profiling of perturbation responses across many cell contexts using single-cell RNA-seq applied to co-treated pools of cancer cell lines. We demonstrate this approach by profiling the responses of pools of 24 - 99 cell lines to a range of different drugs, as well as to CRISPR perturbations.

To demultiplex the data, as well as to identify droplets containing ambient mRNA (empty droplets) or two cells (‘doublets’), we developed a computational method based on SNP-fingerprinting, which classifies single cells with negligible error rates, even for cells with low sequencing depth. During the course of this project, several other methods for SNP-based demultiplexing have been published^20,47,48^. In particular, Demuxlet applies a similar approach, using pre-computed reference SNP profiles for the samples being pooled to identify single cells and detect doublets. Indeed, for data sets where we could run both models, we found that they produced identical single-cell classifications, and similar doublet detection results (**Supplementary Fig. 1**). The key advantage of our approach is that it can be applied to very large pools of samples. We demonstrate this directly on pools of ∼100 cell lines, and our analysis shows that our method can be applied with even larger pools of hundreds of cell lines. This is because we detect doublets by leveraging a Lasso-regularized generalized linear model (Methods) that efficiently estimates the most likely mixture of two reference SNP profiles for each single cell. In contrast, Demuxlet detects doublets by explicitly computing the likelihood of all possible pairwise combinations of samples, making it computationally intractable for such larger pools^20^.

A number of approaches have been developed for high-throughput transcriptional profiling that can be used to study perturbation responses at scale. The Connectivity Map project has utilized a low-cost bead-based assay that measures a reduced set of 1000 ‘landmark’ genes to profile thousands of different perturbation responses^10,12^. More recently, methods such as DRUG-seq^14^ and PLATE-seq^13^ use oligo-tagging of treatment conditions to perform multiplexed RNA-sequencing, greatly reducing library preparation costs. Similar sample-barcoding strategies have also been employed with scRNA-seq^21,43^, allowing for multiplexed profiling across treatment conditions such as time points and drugs. MIX-Seq complements these existing approaches by allowing for multiplexed profiling of perturbation responses across broad panels of heterogeneous cell contexts, without the need for additional experimental barcoding steps. As we demonstrate, MIX-Seq can also be combined with existing sample-barcoding strategies, such as Cell Hashing, to enable dual-multiplexing across treatment conditions and cell contexts.

The single-cell resolution of the data empowers novel biological analyses as well. For example, we show that analysis of perturbation-induced changes in the inferred cell cycle composition across cell lines can provide insights into mechanisms underlying decreased proliferation. Single-cell profiling could also be used to isolate the effect of perturbations on different cell subpopulations, as well as to measure changes to the overall composition of the cell population, which could be a powerful tool when studying more hetergenous samples, and mechanisms of drug resistance.

MIX-Seq’s ability to efficiently profile genome-wide transcriptional responses across a broad panel of cell lines provides several advantages relative to traditional approaches that assess perturbation responses in a small number of representative cell lines. First, profiling across a broad panel of heterogeneous cell lines allows for detection of context-specific responses. For highly selective drugs like dabrafenib and nutlin it’s often critical to profile cell lines with the appropriate genomic context (i.e. TP53 WT, BRAF mutant) in order to see a clear transcriptional response (e.g. **Fig. 2, Fig. 6**). Even among sensitive cell lines, however, there can be substantial response heterogeneity, and profiling many cell lines naturally makes results less sensitive to the particular choice of cell line models under study. For example, we found that responses to the BRAF-inhibitor dabrafenib showed substantial variation, even among the highly sensitive BRAF-mutant melanoma cell lines (**Supplementary Fig. 15**).

By profiling perturbation responses across large panels of well-characterized cell lines, we can also uncover how patterns of transcriptional changes relate to the underlying genomic and functional features of the cells. In particular, pairing MIX-Seq with PRISM^19^, which can measure long-term drug sensitivity across the same panel of cell lines, allows us to dissect the components of transcriptional response associated with decreased cell viability in order to better understand the mechanisms underlying a drug’s fitness effects. For the drugs studied here, we found that viability-related responses were broadly similar across drugs, mostly reflecting a down-regulation of cell-cycle genes and up-regulation of genes involved in translation (**Supplementary Fig. 9**), though transcriptional signatures associated with apoptosis were also observed for some drugs (*e*.*g*. **Supplementary Fig. 4**). The two clear exceptions to this pattern were both inhibitors of anti-apoptotic proteins -- the BCL-2i navitoclax and the MCL1i AZD5591. These drugs did not produce strong and/or selective transcriptional responses. This suggests that compounds which directly induce apoptosis may not elicit a clear transcriptional signature, at least when measured 24 hour post-treatment as done here.

One potential caveat of profiling transcriptional responses in pools of cell lines is that paracrine signaling between cell lines in the pool could affect the measured responses. We found that scRNA-seq profiles at baseline for cells grown in a pooled format were consistently most similar to bulk RNA-seq measurements of the same cell lines grown individually (**Fig. 1c**), suggesting that such paracrine-signaling effects are likely to be modest. Measuring treatment and control conditions within the same pool of cell lines also provides some internal control for baseline effects of paracrine signaling. Finally, previous work has shown that drug response profiles measured in cell line pools are largely concordant with standard measurements^19^. Nevertheless, the potential for interactions between cell lines in the pool must be considered when measuring perturbations responses using MIX-Seq.

We also used MIX-Seq to show that transcriptional responses measured 6 - 24 hours after drug treatment can be used to predict long-term cell viability remarkably well across a handful of targeted cancer drugs. Notably, for the drugs tested here, the ability of machine learning models to predict drug sensitivity (as measured by conventional cytotoxicity assays) from a cell line’s transcriptional response was substantially better than when using baseline ‘omics’ features. These results suggest that transcriptional profiling could be used as a robust functional pharmacodynamic marker of drug sensitivity, which may provide improved predictions of tumor vulnerabilities compared with standard biomarker approaches. An important potential future application of this approach would be to utilize scRNA-seq to rapidly assess the sensitivity of primary tumor cells to various drug treatments *ex vivo*, circumventing the prolonged primary cell cultures needed to achieve sufficient cell numbers for standard long-term viability assays such as Cell-Titer-Glo^49,50^. However, it will be important to extend these tests to a broader range of drugs, and primary patient-derived cancer models, in order to understand the generalizability of these results.

We envision that MIX-Seq could be used to efficiently build a database of transcriptomic changes elicited by a broad range of different chemical and genetic perturbations, each measured across a large heterogeneous panel of cancer models. Analogously to the CMAP project^10,12^, such a database could be used for predicting the mode of action of compounds and genetic manipulations whose cellular effects remain to be uncovered. By measuring perturbation responses across many different cell contexts, with single-cell resolution, MIX-Seq provides a powerful tool for identifying the core transcriptional programs of cancer cells, and better understanding how perturbations interact with the underlying cell context to alter these programs.

## Acknowledgements

This work was funded in part by the Lustgarten Foundation (AJA, BMW), Dana-Farber Cancer Institute Hale Center for Pancreatic Cancer Research (AJA, BMW), the Doris Duke Charitable Foundation (AJA), Pancreatic Cancer Action Network (AJA), NIH-NCI K08 CA218420-02 (AJA), P50CA127003 (AJA) and U01 CA224146 (AJA).

## Author Contributions

Conceptualization and design, JMM, BRP, AW, KG-S, TS, MR, IT, AR, AJA, FV, AT; Experiments, BRP, KG-S, TS, MR, OK, DD, SB; Data analysis, JMM, AW, AJ; Writing, JMM, BRP, AW, AR, AJA, FV, AT; Supervision, OR-R, JAR, TRG, AR, AJA, FV, AT; Project Administration, EC, DD; Funding Acquisition, AJA, BMW, TRG, AR.

## Methods

### Method of cell line pooling

Cell line pools were made in sets of 25 cell lines. These 25 cell line pools were chosen based on doubling time and were grown in RPMI without phenol red and with 10% FBS. Cell lines were then washed with 10 mLs of PBS, and trypsinized with 1 mL trypsin which was then removed. Ten mLs of RPMI media were added to the cells post trypsinization and resuspended. Cells were then counted by a Nexcelom cellometer using 10 uL of cell suspension and 10ul of Trypan blue. Equal numbers of cells per cell line were mixed together and spun down at 1,250 RPM for 5 minutes. Media was aspirated and the cells were resuspended in Sigma Cell Freezing media and frozen in 1 mL aliquots. This process was repeated for all of the 25 cell line pools. For MIX-Seq experiments involving larger pools, multiple 25 cell line pools were thawed in RPMI with 10% FBS, spun down and resuspended in 5 mLs of RPMI media. Cells were then counted and equal numbers were combined together on the day of plating to form larger pools of up to ∼100 cell lines.

### Cell culture

For drug treatment experiments, cell line pools were cultured in RPMI containing 10% fetal bovine serum but did not contain phenol red or penicillin/streptomycin. Cell line pools were validated as mycoplasma free prior to initiating the experiment. Cell line pools were plated at 200,000 cells per well in 6 well plates containing 2 mL of RPMI culture media described above. Cell seeding density did not vary depending on pool size (∼25, 50 or 100 cell line pools). Cell pools were plated ∼16-20 hours prior to drug treatment. Cells were treated with the described drugs or vehicle (DMSO) with a 0.2% final media DMSO concentration.

For GPX4 knockout, cell line pools were plated at 200,000 cells per well in 12 well plates containing 1 mL of RPMI culture media. 24 hours later, the cells were infected with lentivirus expressing Cas9 and sgRNA at a multiplicity of infection of 20 in the presence of 4 ug/mL of polybrene. At 48 hours after the infection, the culture medium was replaced with medium containing 1 ug/mL puromycin. Cells were harvested at 72 or 96 hours after the infection.

### Cell harvesting

Generally cells were harvested after drug treatment using standard cell culture methods. After drug treatment, cells that were in suspension (presumably containing dead cells from drug treatment) were collected and reserved for addition to the adherent cell fraction. Adherent cells were washed once with 1x PBS, trypsinized in 1 mL trypsin, incubated for 3-7 minutes are 37C, and then trypsin inactivated with 1 mL growth media. For cell hashing, cells were treated with TrypLE Express (ThermoFisher) instead of trypsin to reduce the amount of cell surface proteins digested that may affect the binding of cell hashing antibodies.

For the trametinib time course experiment, cells were treated with trametinib with a staggered dosing schedule so all timepoints could be collected simultaneously. Cells were plated 19 hours prior to the first drug treatment, corresponding to the 48 hour time point. Cells were harvested for 10X capture ∼67 hours after initial seeding. Final concentrations for drug treatments are listed in **Supplementary Table 3**.

### Preparation of cell suspensions and scRNA-Seq

After trypsinization, adherent and suspension cells were combined for each treatment, pelleted, and resuspended in Cell Capture Buffer (1x PBS with 0.04% BSA). Cells were counted (including trypan blue non viable cells) and resuspended at a concentration of 1,000 cells per microliter for standard loading on the Chromium Controller (10x Genomics), or at 1,500 cells per microliter for “super loaded” samples. Up to 40,000 cells were loaded per 10x channel for “super loaded” samples, with expected recovery of up to 20,000 cells per channel. Cell suspensions were captured on a 10x Chromium controller using Single Cell 3’ reagent chemistries (either version 2 or version 3 reagents) (**Supplementary Table 2**).

### Cell Hashing cell labelling

Cell Hashing^21^ was performed using the cell harvest method described above with the following changes. All steps were performed on ice. Harvested cells were resuspended in Cell Hashing Staining Buffer (1x PBS with 2% BSA and 0.02% Tween) prior to cell counting. Samples were counted in duplicate with two technical replicates by Countess (Life Technologies) to estimate total cell number. Up to 1,000,000 cells (range 3e5-1e6 cells) were resuspended in 100 microliters of Cell Hashing Staining Buffer. Cells were blocked with 10 microliters of Human TruStain FcX blocking solution (BioLegend) for 10 minutes at 4C. Cells were then incubated with 2µL (1µg) of the appropriate BioLegend TotalSeq™-A Hashing antibody (product codes: 394607, 394609, 394611, 394613, 394615, 394617, 394619, 394623, 394625, 394627, 394629) for 30 minutes at 4C. Cells were washed three times with 0.5mL of Cell Hashing Staining Buffer and filtered through low volume 40µm cell strainers (Flowmi). All cell suspensions were recounted to achieve a uniform concentration of 1,500 cells per microliter before pooling for 10x capture.

### Cell hashing library preparation

Separation of hashtag oligo (HTO)-derived cDNAs (<180bp) and mRNA-derived cDNAs (>300bp) was done after whole transcriptome amplification by performing 0.6X SPRI bead purification (Agencourt) on cDNA reactions as described in 10x Genomics protocol. Briefly, the supernatant from 0.6X SPRI purification contains the HTO fraction, which was subsequently purified using two 2X SPRI purifications per manufacturer protocol (Agencourt). HTO’s were eluted by resuspending SPRI beads in 15 µl TE.

Purified HTO sequencing libraries were then amplified by PCR. PCR reactions were as follows:

| Reagent | Volume |
| --- | --- |
| Purified HTO fraction after 2X SPRI | ~1 uL (5 ng) |
| 2x NEB Next Master Mix | 25 uL |
| Illumina TruSeq DNA D7xx_s primer (containing i7 index) 10 μM | 1 uL |
| SI PCR oligo 10 μM | 1 uL |
| H<sub>2</sub>O | To 50 uL final volume |

Typically 3 identical “dial out” PCR reactions were performed per HTO library. We varied the number of PCR cycles to avoid under or overamplifying the HTO libraries. PCR cycling conditions were are follows

| Step | Temperature | Time |
| --- | --- | --- |
| 1 | 98C | 10 sec |
| 2* | 98C | 2 sec |
| 3* | 72C | 15 sec |
| 4 | 72C | 1 min |

\*Steps 2&3 were repeated for 15, 18, or 22 cycles

PCR reactions were purified using another 2X SPRI clean up and eluted in 15 µL of 1x TE. HTO libraries were then analyzed for amplification quality. Libraries were quantified by Qubit High sensitivity DNA assay (ThermoFisher) and loaded onto a BioAnalyzer high sensitivity DNA chip (Agilent) to determine if an intended HTO product size of ∼180 bp was achieved.

### Sequencing

Samples were sequenced using HiSeq X (Illumina) and NovaSeq 6000 (Illumina) platforms. The read structure (for 10x 3’ v3 chemistry) was as follows:

| Platform | Read | Cycles |
| --- | --- | --- |
| HiSeq | Read 1 | 28 (26)* |
|  | Read 2 | 96 |
|  | Index 1 | 8 |
| NovaSeq | Read 1 | 28 (26)* |
|  | Read 2 | 80 |
|  | Index 1 | 8 |

\*For 10x 3' v2 chemistry

The hashing library for the trametinib time-course experiment was sequenced twice with spike-ins of 2.5-10%.

### Data processing

Sequencing data were processed using 10x Cell Ranger software, run using the ‘Cumulus’ cloud-based analysis framework^51^. Our initial experiments were done with 10x Single Cell 3’ v2 chemistry, and were processed using version 2 of the Cell Ranger software. In our later experiments we used v3 chemistry, and the corresponding version 3 of Cell Ranger. Reads were aligned to the hg19 reference genome.

### SNP identification

To define a SNP panel for cell line classification, we identified SNPs that occurred frequently across a large panel of 1,160 cell lines, and that were also detected in scRNA-seq data. Specifically, we ran MuTect 1 (version 1.1.6) to call SNVs from bulk RNA-seq data and scRNA-seq data from 200 cell lines using a downsample to coverage rate of 1,000 and a fraction contamination rate of 0.02, and with all other parameters set to defaults. We took the subset of SNPs that were observed in both the bulk and single-cell data, then ordered all SNPs by the frequency of their occurrence (in the bulk RNA-seq data), selecting the 100,000 most frequently observed SNPs.

For the bulk RNA-seq data, we used Freebayes^52^ to estimate allelic fractions across the reference SNP panel, using the settings “pooled-continuous” and “report-monomorphic”, and adding a pseudocount of 1 to the reference and alternate allele read counts. For the single-cell data, we used the method scAlleleCount (https://github.com/barkasn/scAlleleCount) to extract reference and alternate allele counts at all SNP sites.

### SNP-based cell line classification

To estimate the likelihood of the observed SNP reads for an individual cell having come from each reference parental cell line, we use a generalized linear modeling approach. Specifically, we use a logistic regression model, where the probability of a read at SNP site *i* being an alternate allele is given by: *π*_*i*_ = *σ* (*β*_0_ + *β*_*j*_ *X*_*ij*_), where *σ* is the logistic function, *X*_*ij*_ is the allelic fraction of the given cell line *j* at site *i* (estimated from bulk RNA-seq data), and *β* are parameters estimated for each single cell and reference cell line by maximizing the likelihood: 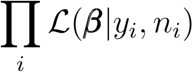, where *ℒ* is the binomial likelihood, *Y*_*i*_ is the number of alternate reads, and *n*_*i*_ is the total number of reads observed at site *i*. We fit models using the glm function in R, and the cell line whose SNP profile X_*j*_ produced the highest likelihood for the observed single-cell SNP reads was selected. Goodness-of-fit was quantified by the deviance ratio: 1 – deviance_fit/deviance_null. We also compute a measure of the classification confidence, given by the margin between the best-fitting and second-best-fitting model deviance ratios, normalized by the standard deviation of deviance ratio values across reference cell lines (excluding the best matching cell line *j*\*).

Estimates of the SNP classification error rate were given by 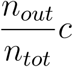, where *n*_*out*_ is the number of cells erroneously classified as out-of-pool cell lines, *n*_*tot*_ is the total number of cells recovered in the experiment (excluding doublets and low-quality cells), and *c* is a correction factor to account for the probability of cells being classified incorrectly among the ‘in-pool’ cell lines. Assuming errors are made with equal probability among all reference cell lines, this is given by 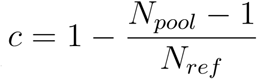.

### Modeling doublets

Doublet detection is performed using a similar generalized linear modeling approach, where alternate allele probabilities are modeled as a mixture of the allelic fraction profiles from two reference cell lines X_*j*_ and X_*k*_: *π*_*i*_ = *σ*(*β*_0_ + *β*_*j*_ *X*_*ij*_ + *β*_*k*_ *X*_*ik*_), where the ratio *β*_*j*_ */ β*_*k*_ represents the proportion of mRNA reads from cell line *j* vs cell line *k*. In order to efficiently estimate the most likely pairwise mixture of reference cell lines, we use a Lasso-regularized generalized linear model (implemented in the R package glmnet^53^), considering the allelic fraction profiles for all in-pool reference cell lines X_*j*_ as covariates. We constrained coefficient estimates to be non-negative, and limiting the model to use a maximum of 2 non-zero coefficients (i.e. 2 reference SNP profiles). After using the Lasso model to estimate the most-likely ‘doublet’ pair of cell lines, we then refit the GLM without regularization to estimate the goodness-of-fit of the doublet model (deviance), as well as the optimal mixing ratio. To measure the evidence in favor of a cell being a doublet, we use the difference of deviance ratios of the best-fit doublet and singlet models (equivalent to the log likelihood ratio of the doublet and singlet models, normalized by the log likelihood ratio between the saturated and null models).

### Classifying low-quality cells and doublets

To identify low-quality cells and classify doublets, we first remove cells which have a high or low proportion of UMIs from mitochondrial genes (> 0.25 or < 0.01), or with reads at fewer than 50 of the reference SNP sites. In many experiments we observed groups of cells with distinct gene expression profiles, and SNP profiles that did not match to any reference cell line (or pairwise combination of cell lines) in particular, but rather resembled more a mixture of SNPs from all the in-pool cell lines, suggesting these were empty droplets containing ambient mRNA in the pool^18,21^. To identify these putative empty droplets, we first clustered the single-cell expression profiles using Seurat’s default graph-based clustering with 10 nearest neighbors and a resolution parameter of 1-4 (depending on the pool size). We then identified gene expression clusters which consistently had poor-fitting SNP models (i.e. that did not resemble singlets or doublets based on their SNPs). For this, the overall SNP model goodness-of-fit for each cell was assessed by the deviance ratio of the doublet model, which was strictly greater than or equal to that of the restricted singlet model. The median SNP-model deviance ratio was computed for each gene-expression cluster, and clusters with a median deviance ratio of less than 0.3 were considered to be low quality, and were removed from the data before further analysis.

We then separated doublets from singlets using a 2-component Gaussian mixture model (GMM) fit with two features: the singlet model deviance ratio, and the doublet model goodness-of-fit (difference in deviance ratios relative to the singlet model). GMMs were fit using the R package MClust^54^, with the default conjugate prior on the covariance matrices, and no shrinkage on the component means. Cells with a probability > 0.5 of being doublets were then taken to be doublets.

Finally, to ensure cells labeled singlets were confidently identified, we also required that the difference in goodness-of-fit between the best-fitting and second best-fitting reference cell lines was at least 2 z-score. Cells that were excluded based on any of the above criteria (other than doublets) were labeled ‘low-quality’ (**Supplmenetary Fig. 1**).

### Visualizing single-cell expression profiles

2D representations of single-cell expression profiles (e.g. **Fig. 1b**) were generated using Seurat v3^55^. Single-cell counts data were first normalized and log-transformed using the NormalizeData function, with a scale_factor of 10^5^. Data were then normalized across cells using the ScaleData function. The top 5,000 most variable genes (based on the ‘vst’ selection method) were selected using the *FindVariableGenes* function, and principal components were computed using the *RunPCA* function, retaining the top 2*N* PCs, where *N* is the number of cell lines in the pool. t-SNE embeddings were computed based on the PCs, using the *RunTSNE* function with a perplexity parameter of 25. UMAP embedings were computed using the *RunUMAP* Seurat function with 15 nearet neighbors, and a ‘min.dist’ parameter of 0.6 (default parameters otherwise).

### Gene-expression based cell line classification

For comparison with SNP-based cell line classification, we also classified single cells based on the similarity of their gene expression profiles to bulk RNA-seq measurements from the parental cell lines (using the 19Q2 DepMap gene expression data; depmap.org)^46^. For this analysis we combined the control datasets for each cell line pool (untreated or DMSO-treated).

Rather than comparing each individual cell’s expression profile with the bulk RNA-seq data directly, we first derived ‘de-noised’ estimates of the single-cell expression profiles by clustering cells and then computing the within-cluster average expression profiles. Specifically, single-cell expression profiles were normalized and scaled, followed by principal component analysis, as described above. We then applied Seurat’s default graph-based clustering with 10 nearest neighbors and a resolution parameter of 1 (24 cell line pool) or 20 (99 cell line pool). After identifying clusters, we sum-collapsed read counts across cells within each cluster, and then transformed the data to log counts-per-million (with a pseudo count parameter of 1). These cluster-averages were taken as estimates of each single cell’s expression profile.

To compare these single-cell profiles with bulk RNA-seq profiles we first mean-centered each dataset across samples per gene. We then identified the 5000 genes (present in bulk and single-cell datasets) with highest variance across bulk RNA-seq samples. Each cell was then classified according to the cell line whose bulk RNA-seq profile was most correlated (Pearson correlation) across these 5,000 genes.

### Differential expression analysis

To estimate the average transcriptional response of each cell line to a perturbation, we first sum-collapsed the data -- summing read counts across cells for each cell line and treatment condition -- to produce a bulk RNA-seq style read counts profile for each sample^56,57^. We then computed normalization factors per sample (cell line and condition) using the “TMMwzp” method from the edgeR R package^58^, and transformed the profiles to log counts-per-million (using a ‘pseudo count’ of 1) using the edgeR function ‘cpm’ before computing the log-fold-change difference in relative gene abundances between treatment and control conditions.

Differential expression analyses across cell lines was performed using the “limma-trend” pipeline^29,30^, applied to these sum-collapsed and normalized profiles. For this analysis we included data from cells both 6 and 24 hours post-treatment with vehicle control (DMSO) in the control group, as we did not observe a consistent time-related effect of DMSO treatment in our data (e.g. **Supplementary Fig. 12**). Global differences between the two control conditions were incorporated into the model to help mitigate batch effects^57^.

To identify the average drug response across cell lines we thus used models of the form:

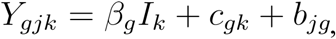

where 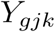, the logCPM expression level of gene *g* in cell line *j* and condition *k*, is modeled as a sum of several terms. The first term captures the average treatment effect, where *β*_*g*_ is the average LFC of gene *g* in response to treatment and *I*_*k*_ is an indicator variable representing whether condition *k* is treatment or control. The second term captures differences in average expression across the control conditions, and the final term captures the baseline expression of each cell line.

To estimate the ‘viability-related’ and viability-independent’ response components, we used a similar modeling approach, including the measured drug sensitivity of each cell line as a covariate interacting with treatment as follows:

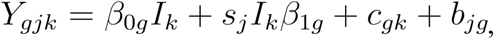

where *s*_*j*_ is the measured sensitivity of cell line *j* to the treatment (one minus the area under the dose-response curve), 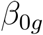 is the ‘viability-independent’ response of gene *g* to treatment, and 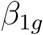 is the viability-related response of gene *g* to treatment.

Only genes with at least 5 reads detected (summed across cells) in at least 5% of the samples were included in analysis. p-values were derived from empirical-Bayes moderated t-statistics, and FDR-adjusted p-values were obtained using the Benjamini-Hochberg method^59^.

When comparing the transcriptional responses of two cell lines to a drug (*e*.*g*. **Supplementary Fig. 14**), the above method cannot be applied as there would be a single sample for each condition. Hence, we compared the uncollapsed single-cell expression profiles. Specifically, we used the edgeR quasi-likelihood approach^60^, following the pipeline used in^61^, including cell detection rate (the fraction of genes with non-zero reads detected) as a covariate.

### Drug sensitivity data

Cell line drug sensitivity data were taken from the Sanger GDSC dataset^4,7^, as well as data generated using the PRISM multiplexed drug screening platform^16,19^. For most compounds we used the area under the dose-response curve (AUC) to measure sensitivity. When data were available from both PRISM and GDSC datasets for a given drug, we used the average of each cell line’s AUC values, after quantile normalization of the AUC measurements from each dataset.

For nutlin treatment, we combined nutlin-3a data from GDSC with PRISM data for the nutlin-family compound idasanutlin (RG7388). For the tool compound BRD-3379, we found that the (PRISM) data were most reliable for the highest dose, so we used log viability measurements at a single dose of 10 uM, though results were similar when using the AUC.

### Gene set enrichment analysis

For analysis of gene set enrichment of transcriptional response signatures, we used a simple approach, measuring the set overlap (Fisher’s-exact test) between each gene set and the 50 top up- and down-regulated genes across (based on the estimated log-fold-change). The collection of gene sets used was the combination of the ‘Hallmark’, and ‘Canonical’ gene set collections from MSigDB v6.2^62^.

### Estimating relative cell line abundance

Estimates of the effects of perturbations on relative cell line abundance were obtained by counting the number of (QC-passing) single cells from each cell line in each treatment condition, adding a ‘pseudo-count’ of 1, and normalizing counts across cell lines per condition. These relative abundance estimates were averaged across samples for each treatment condition to compute the log_2_-fold-change difference between drug-treated and control relative cell line abundances.

### Cell cycle analysis

Cell cycle phase classification was performed with the Seurat function CellCycleScoring, using the S- and G2M-phase gene lists reported in^31^. The change in proportion of cells in each phase between treatment and control conditions, along with associated confidence intervals, were estimated using the prop.test R function for each cell line. For **Fig. 3c**, we computed aggregate scores representing how each compound altered the cell cycle composition by computing weighted averages across cell lines of the change in proportion of cells in each phase, where the weights were determined by the cell lines’ measured drug sensitivity (1 minus AUC, bounded between 0 and 1).

### Principal component analysis

For PCA (and other machine learning analyses), we used a slightly different procedure to estimate each cell line’s average transcriptional response to drug treatment. Rather than ‘sum-collapsing’ the read count data, we ‘mean-collapsed’ the single-cell gene expression profiles by normalizing each single-cell profile to counts-per-million, averaging across cells, and then log-transforming the averaged profiles (using a larger pseudo count value of 10 to help stabilize log-fold change estimates for lowly expressed genes). PCA was then computed on the matrix of cell line log-fold change profiles, mean-centered per gene, using the 5000 genes with most across-cell-line variance. We only used cell lines where there were at least 5 cells in both control and treatment conditions.

The use of mean-collapsed, rather than sum-collapsed, profiles for machine learning analysis helped prevent any bias in the estimated log-fold change responses related to the number of cells recovered for each cell line. Both sum-collapsed and mean-collapsed log-fold change estimates produced similar results, differing primarily in whether cells with greater sequencing depth are given more weight.

Comparisons of PC1 loadings with measured drug sensitivity across cell lines (**Supplementary Fig. 13**) were made using Pearson correlations, with p-values estimated using the ‘cor.test’ R function. FDR adjusted p-values were estimated using the Benjamini-Hochberg method^59^.

### Transcriptional response embedding

To compute the embedding of transcriptional response profiles (**Fig. 6f**), we used the UMAP method^45^, as implemented in the Seurat package. Specifically, we compiled all log-fold change response profiles across cell lines and treatment conditions (computed using mean-collapsed profiles). We restricted analysis to response profiles supported with at least 10 cells per condition and 40 cells total. We then took the 5,000 genes with highest variance across the selected profiles, and computed the top 30 principal components. UMAP was then run using cosine distance between samples in this principle component space, with an ‘n.neighbors’ parameter of 15, and ‘min.dist’ of 0.6.

### Predictive modeling analysis

To assess how well we could predict a cell line’s drug sensitivity from baseline features, or measured transcriptomic responses, we used random forest regression models (implemented in the R package *ranger*^63^) with default parameters. Prediction accuracy (R^2^ of model predictions) was evaluated using 10-fold cross-validation. To help mitigate overfitting, we also applied a pre-filtering of the features, selecting the top 1000 features based on the magnitude of their marginal correlation with the response variable (feature selection was performed separately for each cross-validation set, using training data only).

For the ‘baseline omics’ features, we used baseline logTPM expression levels of each protein coding gene, as well as the damaging and hotspot missense mutation status of each gene. These data were taken from the DepMap 19Q2 data release, available at depmap.org^6,46^.

We only included cell lines with at least 5 cells per condition (treatment and control) for a given drug to ensure the estimated transcriptional response profiles were sufficiently robust.

### Time course analysis

Classification of single-cell treatment conditions, as well as doublet classification, from the hashtag read counts data was performed using DemuxEM^64^, with default parameters.

We used the same approach described above to estimate the viability-related and viability-independent components of the response at each time point post-trametinib treatment. Since we did not observe substantial transcriptional changes across time points after DMSO-treatment (**Supplementary Fig. 10**), we pooled together data across DMSO conditions for analysis.

For **figure 5e,f**, we plotted the time course of viability-independent and viability-related responses for the top 10 down-regulated genes in each component, taking the coefficient with the largest magnitude across post-treatment time points for each gene (after filtering for coefficients with FDR < 0.1).

### Sub-population heterogeneity analysis

Analysis of cell subpopulations for a given cell line were performed by first restricting analysis to cells from the target cell line, then using the scTransform method to normalize the data and identify variable genes (we used 5,000 genes)^65^. We then used Seurat’s default clustering methods to identify sub-populations of cells (with 10 PCs, 20 nearest neighbors, and a clustering resolution parameter of 0.25). When identifying subpopulations across experimental conditions (treatment and control), we first regressed out experimental condition as a covariate (using the ‘vars.to.regress’ input of the SCTransform function) before applying the above clustering procedure.

### GPX4 analysis

Differential expression analysis of GPX4 KO was done by comparing the average effects of the two GPX4 targeting guides against the average of the two control guides (one targeting and one non-targeting), following the same analysis procedure as used for drug treatment data. We identified GPX4 dependent and non-dependent lines using the estimated probability of GPX4 dependency for each cell line from the Achilles 19q2 “gene dependency” file (DepMap, Broad, 2019). Cell lines with GPX4 dependency probability greater than 0.5 were considered dependent.

## Data and code availability

All data reported in this manuscript, including single-cell RNA-sequencing data, drug sensitivity measures, and other cell line features used in the analysis, can be accessed at https://figshare.com/articles/MIX-seq_data/10298696.

Custom code used in the analysis, and for generating all figures, is available at https://github.com/broadinstitute/mix_seq_ms. Code used for SNP classification is available at https://github.com/acwarren/single_cell_classification.

## References

1. Tsherniak, A. et al. Defining a cancer dependency map. Cell 170, 564–576.e16 (2017).

2. Meyers, R. M. et al. Computational correction of copy number effect improves specificity of CRISPR-Cas9 essentiality screens in cancer cells. Nat. Genet. 49, 1779–1784 (2017).

3. McDonald, E. R. et al. Project DRIVE: A Compendium of Cancer Dependencies and Synthetic Lethal Relationships Uncovered by Large-Scale, Deep RNAi Screening. Cell 170, 577–592.e10 (2017).

4. Iorio, F. et al. A landscape of pharmacogenomic interactions in cancer. Cell 166, 740–754 (2016).

5. Behan, F. M. et al. Prioritization of cancer therapeutic targets using CRISPR-Cas9 screens. Nature 568, 511–516 (2019).

6. Barretina, J. et al. The Cancer Cell Line Encyclopedia enables predictive modelling of anticancer drug sensitivity. Nature 483, 603–607 (2012).

7. Garnett, M. J. et al. Systematic identification of genomic markers of drug sensitivity in cancer cells. Nature 483, 570–575 (2012).

8. Dixit, A. et al. Perturb-Seq: Dissecting Molecular Circuits with Scalable Single-Cell RNA Profiling of Pooled Genetic Screens. Cell 167, 1853–1866.e17 (2016).

9. Adamson, B. et al. A Multiplexed Single-Cell CRISPR Screening Platform Enables Systematic Dissection of the Unfolded Protein Response. Cell 167, 1867–1882.e21 (2016).

10. Subramanian, A. et al. A next generation connectivity map: L1000 platform and the first 1,000,000 profiles. Cell 171, 1437–1452.e17 (2017).

11. Norman, T. M. et al. Exploring genetic interaction manifolds constructed from rich phenotypes. BioRxiv (2019). doi:10.1101/601096

12. Lamb, J. et al. The Connectivity Map: using gene-expression signatures to connect small molecules, genes, and disease. Science 313, 1929–1935 (2006).

13. Bush, E. C. et al. PLATE-Seq for genome-wide regulatory network analysis of high-throughput screens. Nat. Commun. 8, 105 (2017).

14. Ye, C. et al. DRUG-seq for miniaturized high-throughput transcriptome profiling in drug discovery. Nat. Commun. 9, 4307 (2018).

15. Basu, A. et al. An interactive resource to identify cancer genetic and lineage dependencies targeted by small molecules. Cell 154, 1151–1161 (2013).

16. Corsello, S. M. et al. Non-oncology drugs are a source of previously unappreciated anti-cancer activity. BioRxiv (2019). doi:10.1101/730119

17. Klein, A. M. et al. Droplet barcoding for single-cell transcriptomics applied to embryonic stem cells. Cell 161, 1187–1201 (2015).

18. Macosko, E. Z. et al. Highly Parallel Genome-wide Expression Profiling of Individual Cells Using Nanoliter Droplets. Cell 161, 1202–1214 (2015).

19. Yu, C. et al. High-throughput identification of genotype-specific cancer vulnerabilities in mixtures of barcoded tumor cell lines. Nat. Biotechnol. 34, 419–423 (2016).

20. Kang, H. M. et al. Multiplexed droplet single-cell RNA-sequencing using natural genetic variation. Nat. Biotechnol. 36, 89–94 (2018).

21. Stoeckius, M. et al. Cell Hashing with barcoded antibodies enables multiplexing and doublet detection for single cell genomics. Genome Biol. 19, 224 (2018).

22. Zheng, G. X. Y. et al. Massively parallel digital transcriptional profiling of single cells. Nat. Commun. 8, 14049 (2017).

23. Vassilev, L. T. et al. In vivo activation of the p53 pathway by small-molecule antagonists of MDM2. Science 303, 844–848 (2004).

24. Lamb, J. et al. A mechanism of cyclin D1 action encoded in the patterns of gene expression in human cancer. Cell 114, 323–334 (2003).

25. Lamb, J. The Connectivity Map: a new tool for biomedical research. Nat. Rev. Cancer 7, 54–60 (2007).

26. Viswanathan, V. S. et al. Dependency of a therapy-resistant state of cancer cells on a lipid peroxidase pathway. Nature 547, 453–457 (2017).

27. Jeganathan, S. & Lee, J. M. Binding of elongation factor eEF1A2 to phosphatidylinositol 4-kinase beta stimulates lipid kinase activity and phosphatidylinositol 4-phosphate generation. J. Biol. Chem. 282, 372–380 (2007).

28. Calle, Y., Palomares, T., Castro, B., del Olmo, M. & Alonso-Varona, A. Removal of N-glycans from cell surface proteins induces apoptosis by reducing intracellular glutathione levels in the rhabdomyosarcoma cell line S4MH. Biol. Cell 92, 639–646 (2000).

29. Law, C. W., Chen, Y., Shi, W. & Smyth, G. K. voom: Precision weights unlock linear model analysis tools for RNA-seq read counts. Genome Biol. 15, R29 (2014).

30. Ritchie, M. E. et al. limma powers differential expression analyses for RNA-sequencing and microarray studies. Nucleic Acids Res. 43, e47 (2015).

31. Tirosh, I. et al. Dissecting the multicellular ecosystem of metastatic melanoma by single-cell RNA-seq. Science 352, 189–196 (2016).

32. King, C. et al. LY2606368 Causes Replication Catastrophe and Antitumor Effects through CHK1-Dependent Mechanisms. Mol. Cancer Ther. 14, 2004–2013 (2015).

33. Roschke, A. V. et al. Karyotypic complexity of the NCI-60 drug-screening panel. Cancer Res. 63, 8634–8647 (2003).

34. Nagai, Y. et al. Genetic heterogeneity of the epidermal growth factor receptor in non-small cell lung cancer cell lines revealed by a rapid and sensitive detection system, the peptide nucleic acid-locked nucleic acid PCR clamp. Cancer Res. 65, 7276–7282 (2005).

35. Ben-David, U. et al. Genetic and transcriptional evolution alters cancer cell line drug response. Nature 560, 325–330 (2018).

36. Shaffer, S. M. et al. Rare cell variability and drug-induced reprogramming as a mode of cancer drug resistance. Nature 546, 431–435 (2017).

37. Charafe-Jauffret, E. et al. Breast cancer cell lines contain functional cancer stem cells with metastatic capacity and a distinct molecular signature. Cancer Res. 69, 1302–1313 (2009).

38. Jerby-Arnon, L. et al. A cancer cell program promotes T cell exclusion and resistance to checkpoint blockade. Cell 175, 984–997.e24 (2018).

39. Kinker, G. S. et al. Pan-cancer single cell RNA-seq uncovers recurring programs of cellular heterogeneity. BioRxiv (2019). doi:10.1101/807552

40. Shi-Lin, D., Yuan, X., Zhan, S., Luo-Jia, T. & Chao-Yang, T. Trametinib, a novel MEK kinase inhibitor, suppresses lipopolysaccharide-induced tumor necrosis factor (TNF)-α production and endotoxin shock. Biochem. Biophys. Res. Commun. 458, 667–673 (2015).

41. Lulli, D., Carbone, M. L. & Pastore, S. The MEK inhibitors trametinib and cobimetinib induce a type I interferon response in human keratinocytes. Int. J. Mol. Sci. 18, (2017).

42. Sivan, G. & Elroy-Stein, O. Regulation of mRNA Translation during cellular division. Cell Cycle 7, 741–744 (2008).

43. Shin, D., Lee, W., Lee, J. H. & Bang, D. Multiplexed single-cell RNA-seq via transient barcoding for simultaneous expression profiling of various drug perturbations. Sci. Adv. 5, eaav2249 (2019).

44. Lim, C. P., Jain, N. & Cao, X. Stress-induced immediate-early gene, egr-1, involves activation of p38/JNK1. Oncogene 16, 2915–2926 (1998).

45. McInnes, L., Healy, J. & Melville, J. UMAP: Uniform Manifold Approximation and Projection for Dimension Reduction. arXiv (2018).

46. Ghandi, M. et al. Next-generation characterization of the Cancer Cell Line Encyclopedia. Nature 569, 503–508 (2019).

47. Xu, J., Falconer, C. & Coin, L. Genotype-free demultiplexing of pooled single-cell RNA-seq. BioRxiv (2019). doi:10.1101/570614

48. Huang, Y., McCarthy, D. J. & Stegle, O. Vireo: Bayesian demultiplexing of pooled single-cell RNA-seq data without genotype reference. BioRxiv (2019). doi:10.1101/598748

49. Kodack, D. P. et al. Primary Patient-Derived Cancer Cells and Their Potential for Personalized Cancer Patient Care. Cell Rep. 21, 3298–3309 (2017).

50. Tseng, Y.-Y. & Boehm, J. S. From cell lines to living biosensors: new opportunities to prioritize cancer dependencies using ex vivo tumor cultures. Curr. Opin. Genet. Dev. 54, 33–40 (2019).

51. Li, B. et al. Cumulus: a cloud-based data analysis framework for large-scale single-cell and single-nucleus RNA-seq. BioRxiv (2019). doi:10.1101/823682

52. Garrison, E. & Marth, G. Haplotype-based variant detection from short-read sequencing. arXiv (2012).

53. Friedman, J., Hastie, T. & Tibshirani, R. Regularization Paths for Generalized Linear Models via Coordinate Descent. J Stat Softw 33, 1–22 (2010).

54. Scrucca, L., Fop, M., Murphy, T. B. & Raftery, A. E. mclust 5: Clustering, Classification and Density Estimation Using Gaussian Finite Mixture Models. R J. 8, 289–317 (2016).

55. Butler, A., Hoffman, P., Smibert, P., Papalexi, E. & Satija, R. Integrating single-cell transcriptomic data across different conditions, technologies, and species. Nat. Biotechnol. 36, 411–420 (2018).

56. Crowell, H. L. et al. On the discovery of population-specific state transitions from multi-sample multi-condition single-cell RNA sequencing data. bioRxiv (2019).

57. Lun, A. T. L. & Marioni, J. C. Overcoming confounding plate effects in differential expression analyses of single-cell RNA-seq data. Biostatistics 18, 451–464 (2017).

58. Robinson, M. D., McCarthy, D. J. & Smyth, G. K. edgeR: a Bioconductor package for differential expression analysis of digital gene expression data. Bioinformatics 26, 139–140 (2010).

59. Benjamini, Y. & Hochberg, Y. Controlling the false discovery rate: A practical and powerful approach to multiple testing. Journal of the Royal Statistical Society: Series B (Methodological) 57, 289–300 (1995).

60. Lun, A. T. L., Chen, Y. & Smyth, G. K. It’s DE-licious: A Recipe for Differential Expression Analyses of RNA-seq Experiments Using Quasi-Likelihood Methods in edgeR. Methods Mol. Biol. 1418, 391–416 (2016).

61. Soneson, C. & Robinson, M. D. Bias, robustness and scalability in single-cell differential expression analysis. Nat. Methods (2018).

62. Liberzon, A. et al. Molecular signatures database (MSigDB) 3.0. Bioinformatics 27, 1739–1740 (2011).

63. Janitza, S., Celik, E. & Boulesteix, A.-L. A computationally fast variable importance test for random forests for high-dimensional data. Adv. Data Anal. Classif. (2016). doi:10.1007/s11634-016-0276-4

64. Gaublomme, J. T. et al. Nuclei multiplexing with barcoded antibodies for single-nucleus genomics. Nat. Commun. 10, 2907 (2019).

65. Hafemeister, C. & Satija, R. Normalization and variance stabilization of single-cell RNA-seq data using regularized negative binomial regression. BioRxiv (2019). doi:10.1101/576827

