## Supplementary Material for "Multiplexed single-cell profiling of post-perturbation transcriptional responses to define cancer vulnerabilities and therapeutic mechanism of action"

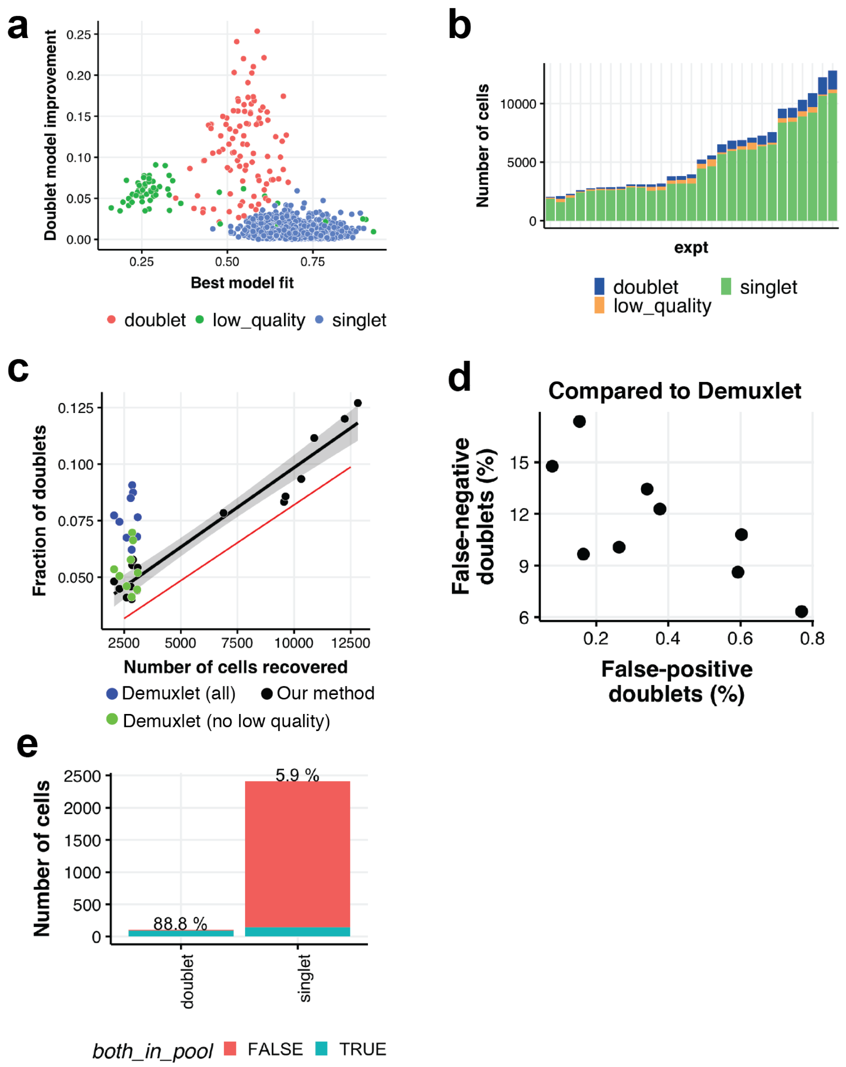


**Supplementary Fig. 1: SNP-based detection of doublets and low-quality cells.**

**a)** Scatterplot of parameters from the SNP-based models (see Methods) from an example experiment, showing classification of cells as singlets, doublets, and 'low-quality'. The y-axis shows the improvement of the doublet model fit over the singlet model, while the x-axis shows the best model goodness-of-fit. **b)** Distribution of cells classified in each category across experiments. **c)** Proportion of cells classified as doublets in each experiment (excluding low-quality cells), as a function of the number of cells recovered in the experiment. Red line shows the trend-line from (Zheng et al., 2017) depicting the expected relationship of doublet probability with cell density. Green and blue points show doublet proportions estimated using Demuxlet (Kang et al., 2018) (only for experiments with smaller pools), with and without exclusion of low-quality cells (based on our analysis pipeline) respectively. **d)** Using the Demuxlet doublet detection as 'ground truth', false-negative and false-positive rates for our doublet classification procedure are shown for each experiment. Overall, Demuxlet tended to produce a somewhat higher rate of detected doublets, even though both methods tended to call doublets at a higher rate than expected based on the cell loading density (**c**). **e)** For an example experiment, we estimated the proportion of cells where the most likely *doublet pair* of reference cell lines were both among the 'in-pool' cell lines (24/494 possible cell lines). 89% of cells classified as doublets had both identified reference cell lines among those in the experimental pool. For cells classified as singlets, there were only ~6% (approximately chance level).

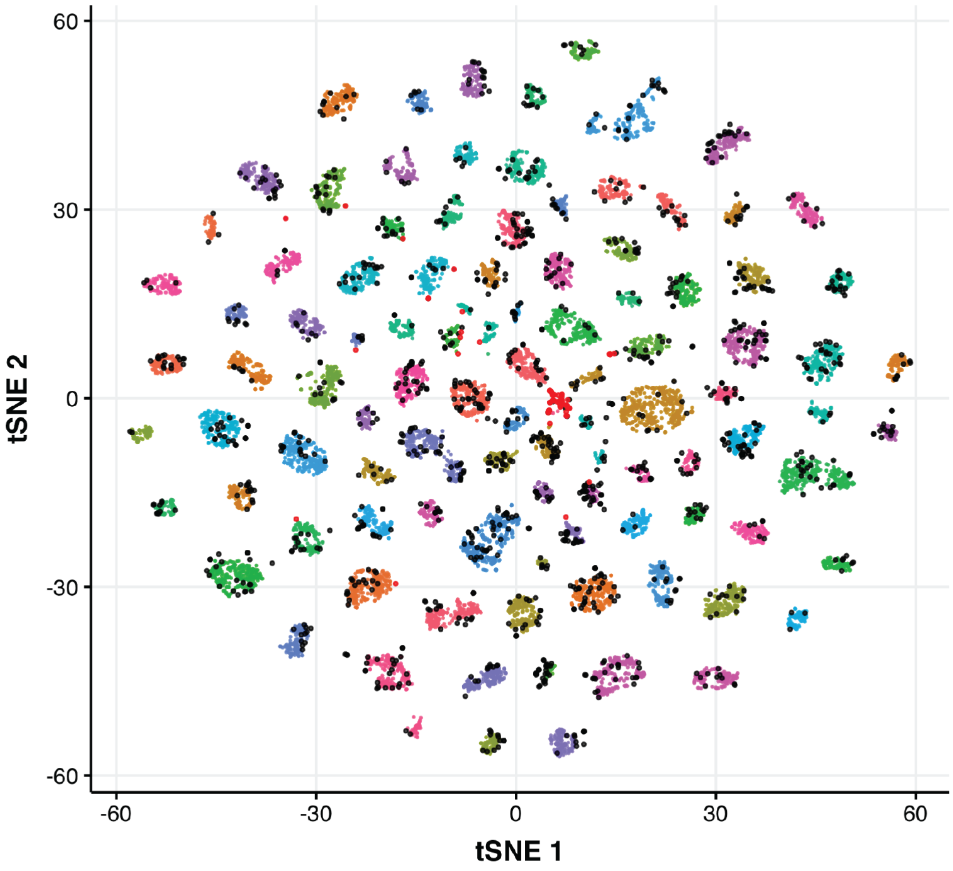


**Supplementary Fig. 2: Agreement between SNP-based and GE-based cell classification.**
Same as **Fig. 1b**, for an example (DMSO-treated) dataset from a 99 cell-line pool. Black dots show cells detected as doublets. Red dots indicate cells where gene expression and SNP-based classifications disagree (0.05% of cells).

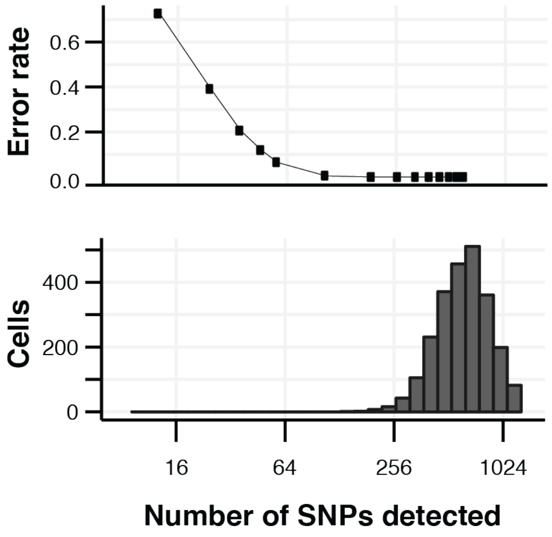


**Supplementary Fig. 3: Dependence of SNP-based cell classification on number of detected SNPs**

(*Top*) Error rate of cell classification was estimated based on the fraction of cells classified among those in the experimental pool (24/494 reference cell lines from this example dataset). Classification accuracy for cells with fewer SNP sites detected was estimated by randomly down-sampling the single-cell SNP reads. (*Bottom*) Distribution of the number of SNP sites detected for the cells measured in the example experiment.

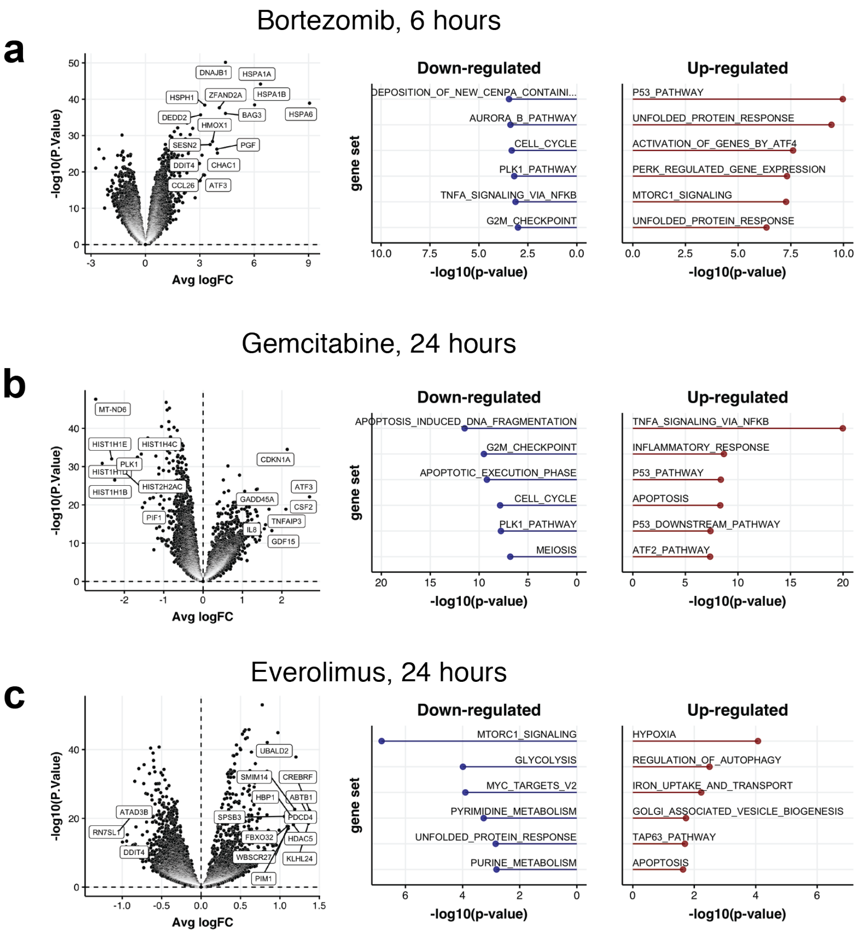


**Supplementary Fig. 4: Identification of compound MOA from transcriptional response profiles.**

**a)** (Left) Volcano plot of the (across-cell-line) average transcriptional response to bortezomib (6 hours post-treatment). (Right) Top gene sets enriched among the most strongly up- and down-regulated genes. **b)** Same as a for gemcitabine treatment (24 hours post-treatment). **c)** Same as **a-b**, for everolimus treatment (24 hours post-treatment).

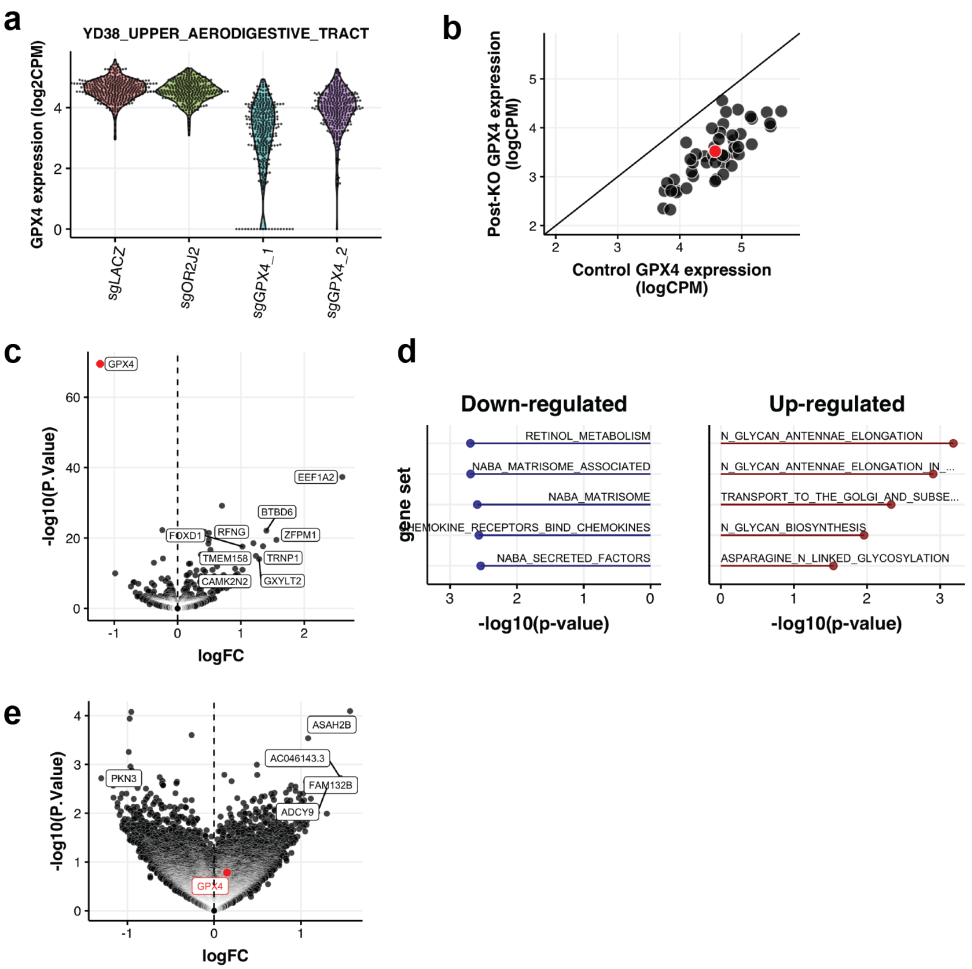


**Supplementary Fig. 5: Pooled transcriptional profiling of genetic perturbation.**

**a)** Distribution of expression levels of GPX4 in an example cell line for cells infected with two sgRNAs targeting GPX4, vs. two control sgRNAs (sgLACZ: non-targeting control, sgOR2J2: 'cutting control'), showing robust on-target knockdown of GPX4 expression. **b)** Comparison of average GPX4 expression across pool of 50 cell lines following GPX4 KO compared with control, showing consistent on-target KD across cell lines. Red dot shows example cell line from **a**. **c)** Volcano plot showing average transcriptional response to GPX4 KO across all cell lines. **d)** Gene set analysis showing up-regulation of glycan related pathways in the average GPX4 KO response. **e)** Volcano plot comparing response to GPX4 KO in GPX4-dependent (n=18) vs. non-dependent (n=15) cell lines. No genes were significantly differentially expressed between the groups.

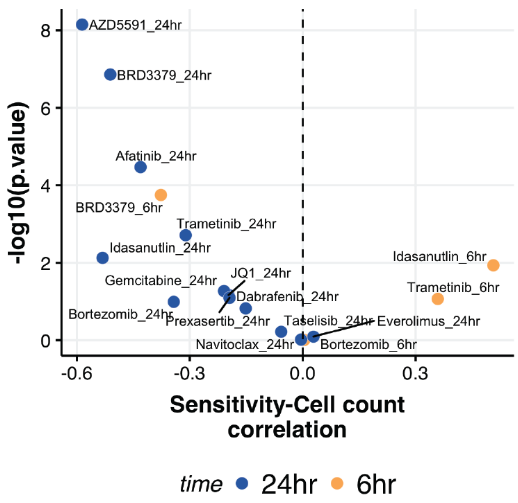


**Supplementary Fig. 6: Direct measurement of relative drug sensitivity via scRNA-seq.** Correlation, and associated p-values, between existing drug sensitivity data (Iorio et al. 2016; Corsello et al. 2019) and the measured change in relative cell abundance (based on pooled scRNA-seq data). The time point of scRNA-seq profiling is indicated by dot color.

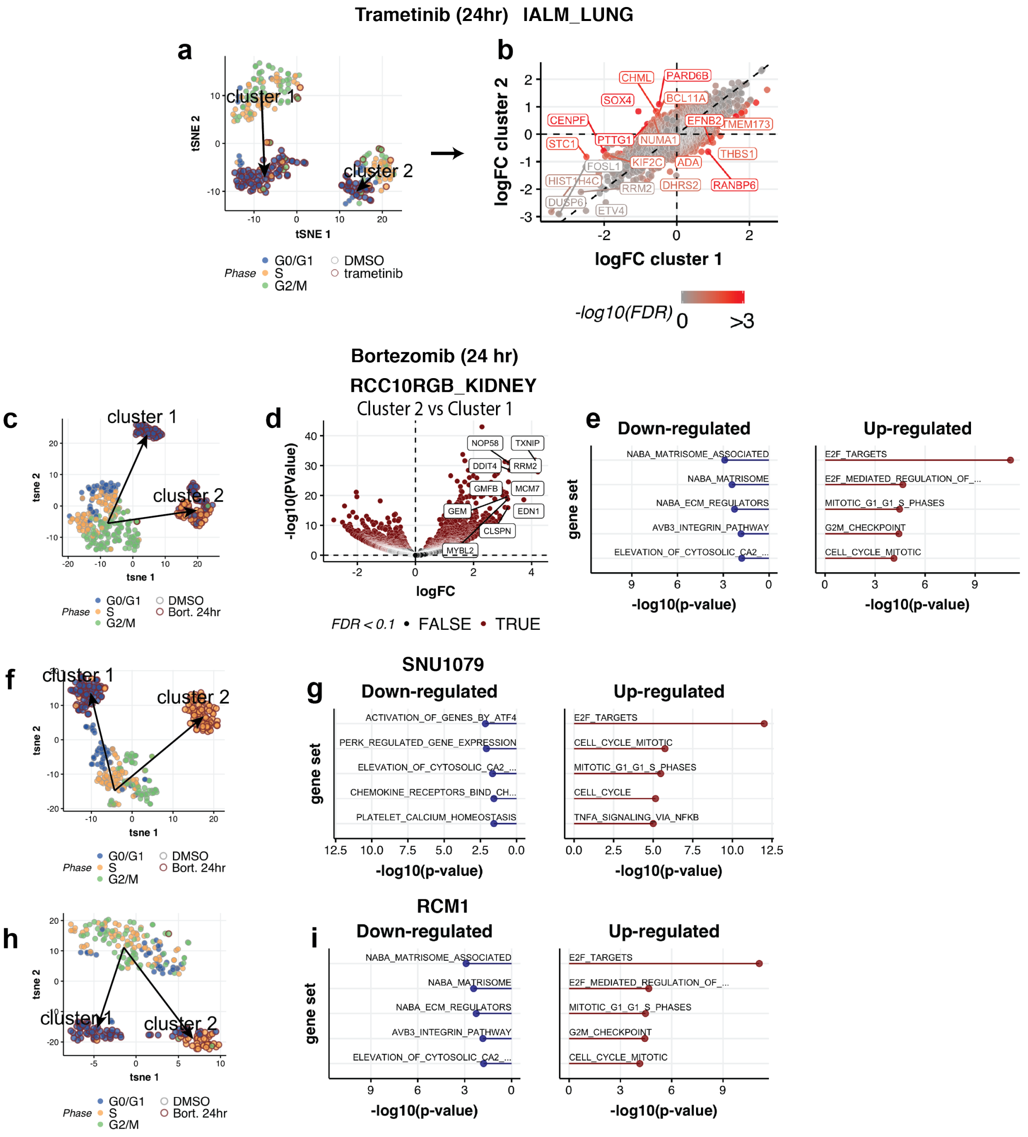


**Supplementary Fig. 7: Single-cell RNA-seq allows analysis of population heterogeneity in drug responses.**

**a)** t-SNE plot showing two distinct sub-populations of IALM cells, and the response to trametinib in each (same as **Fig. 3e**). Fill color depicts inferred cell-cycle phase, and stroke-color depicts treatment condition. **b)** Comparison of the average trametinib response for cells from the two IALM clusters shown in **a**. Dot color represents the significance of the difference between trametinib responses of the two sub-populations. **c)** Same as **Fig. 3f**. **d)** Volcano plot comparing the average expression of the two identified clusters of bortezomib-treated RCC10RGB cells. **e)** Gene set enrichment analysis of the top differentially expressed genes in **d**. **f-i)** Same as **c** and **e**, for the cell lines SNU1079 and RCM1, showing similar bimodal split of the cell populations after bortezomib treatment, with clusters distinguished by expression of E2F targets and cell cycle genes.

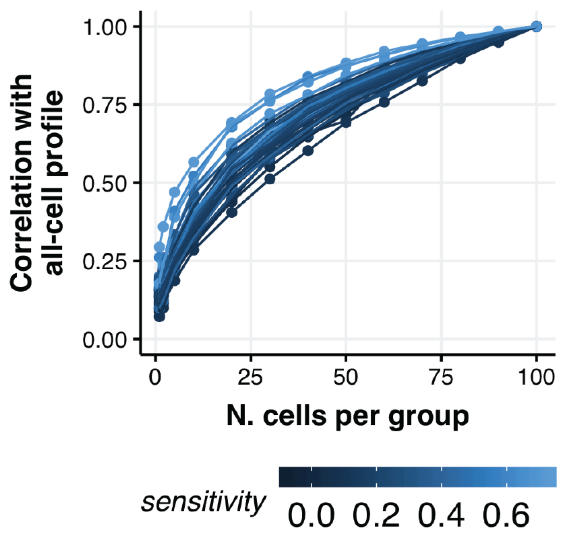


**Supplementary Fig. 8: Impact of cell population size on estimates of transcriptional response profiles.**

To understand how estimates of each cell line's transcriptional response depended on the number of cells sampled per treatment condition we performed a downsampling analysis. Specifically, for measured trametinib response (24 hours post-treatment, as in **Fig. 4**), we took the set of cell lines which had at least 100 cells sampled in each condition (n = 45), and restricted the data to a random set of 100 cells per condition. We then estimated the LFC transcriptional response profile of each cell line using random subsets of cells and compared these subsampled estimates with the profiles derived from the starting set of 100 cells per condition. Profile similarity was assessed by the Pearson correlation of LFC vectors across the 5000 most variably-expressed genes, averaged across 5 repetitions of the downsampling procedure for each cell line. Each line represents data from a different cell line, colored by the cell line's measured trametinib sensitivity (1 – AUC).

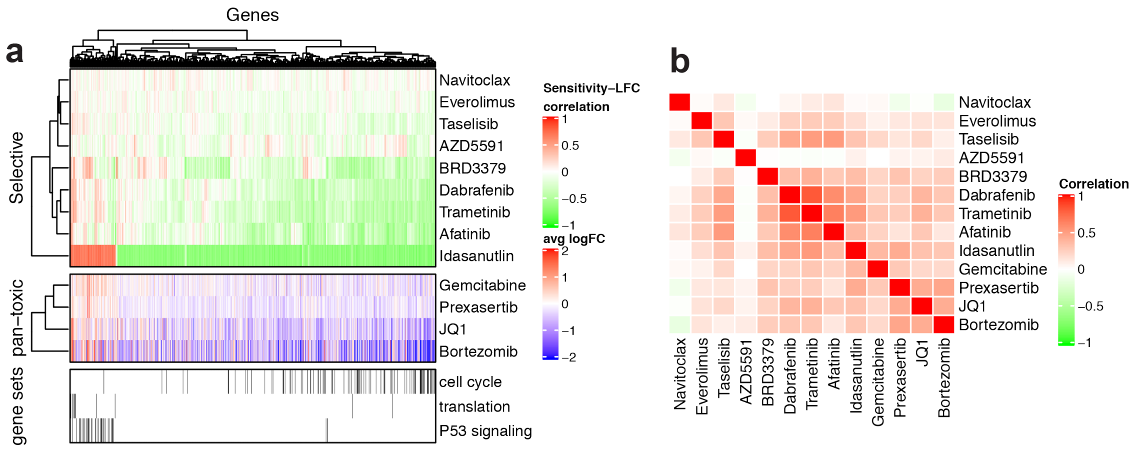


**Supplementary Fig. 9: Similarity of viability-related transcriptional responses across drugs.** **a)** (Top) Heatmap of correlation between transcriptional response and drug sensitivity across cell lines for the 1000 most genes with strongest average correlation for thee 8 selective drugs. (Middle) Heatmap of average logFC drug response across cell lines for 4 pan-toxic drugs tested showing similarity to the viability-related response profiles above. (Bottom) Depiction of gene membership in 3 gene sets (cell cycle = HALLMARK_G2M_CHECKPOINT; translation = REACTOME_TRANSLATION, and P53 signaling = HALLMARK_P53_PATHWAY). **b)** Matrix of correlations for the transcriptional response profiles shown in **a** across all pairs of compounds. Viability-related responses of selective compounds, and average responses for pan-toxic compounds, were broadly similar across these 1000 genes (with the exception of navitoclax and AZD5591).

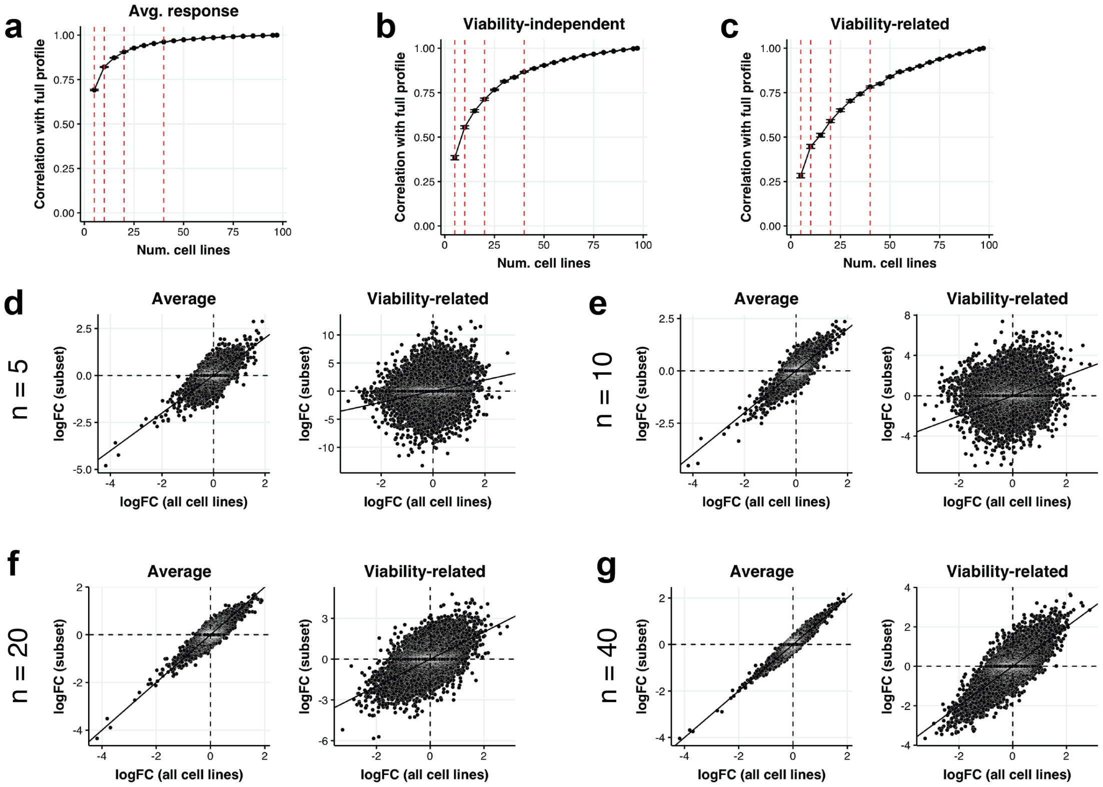


### Supplementary Fig. 10: Effect of cell line sample size on estimation of transcriptional response components.

Analyses of the trametinib response components (as in **Fig. 4**) were repeated with random subsets of the cell lines. **a-c)** Average correlation between the estimated response profile (logFC) when using a subsample of cell lines vs. all the cell lines for the average (**a**), viability-related (**b**), and viability-independent (**c**) components respectively. Error bars show interval +/- s.e.m. Vertical red lines indicate the subsample sizes shown in **d-g**. **d-g)** Scatterplot comparisons of example estimates of average and viability-related response components using all cell lines vs. random subsets of increasing size, from 5 to 40.

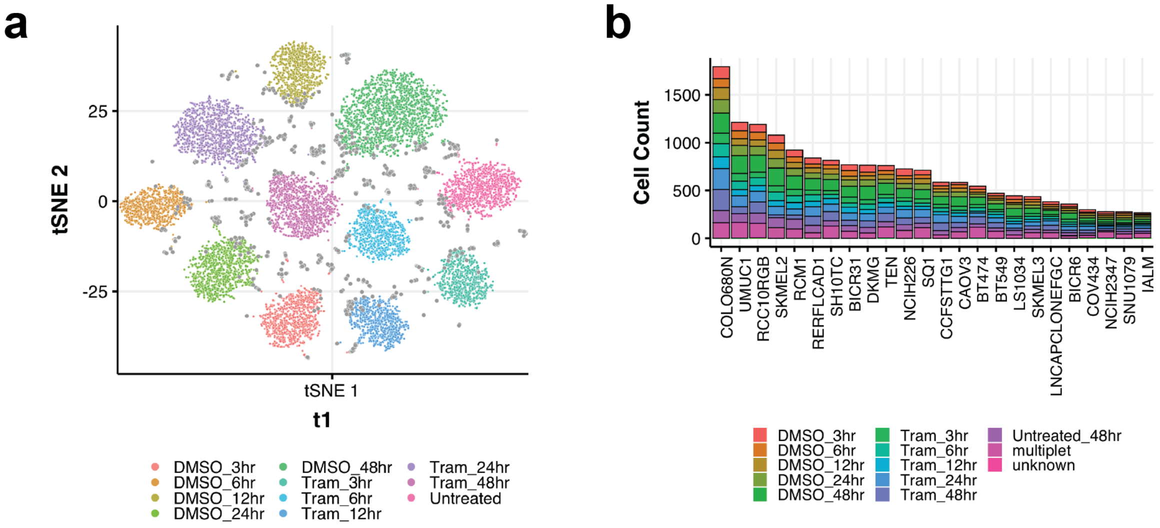


**Supplementary Fig. 11: Cell Hashing provides efficient labeling of cells from different experimental conditions**

**a)** t-SNE representation of hashtag read profiles across single cells from all cell lines, colored by the classified treatment condition. Read count profiles across hashtags were normalized for each cell using the centered log-ratio transformation (with a pseudocount value of 1) prior to computing the t-SNE embedding. Gray dots indicate cells classified as doublets. **b)** Histogram of cell counts by parental cell line and the inferred hashtag condition.

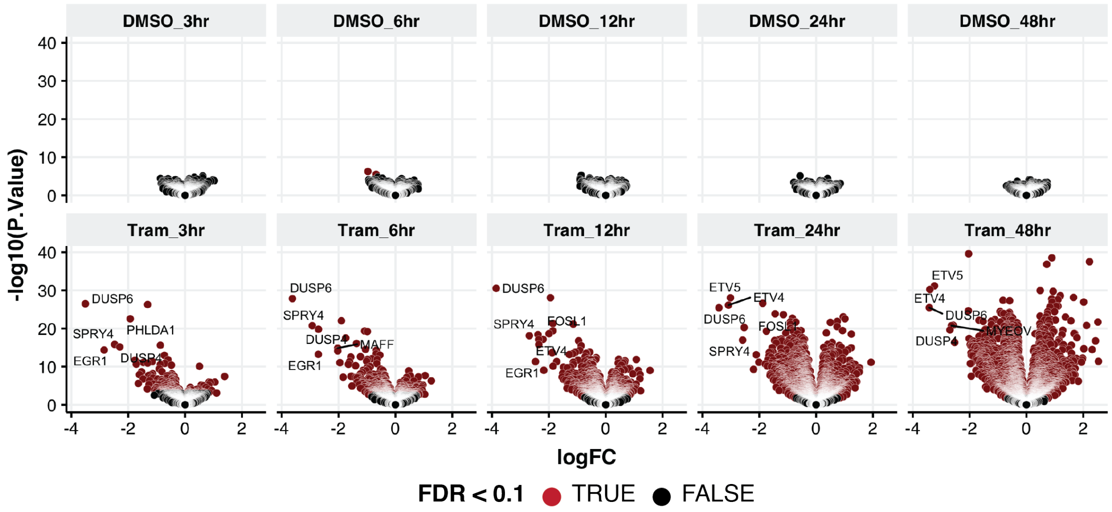


**Supplementary Fig. 12: Lack of time-dependent transcriptional response following treatment with DMSO vehicle control.**

Volcano plots showing the average (across cell lines) transcriptional response to each treatment condition, using untreated cells as reference. Virtually no genes showed significant changes (FDR < 0.1) in any DMSO-treated conditions, and there were no time-dependent trends apparent in the DMSO response. As a result, we combined DMSO conditions, and untreated cells, as reference for other analyses.

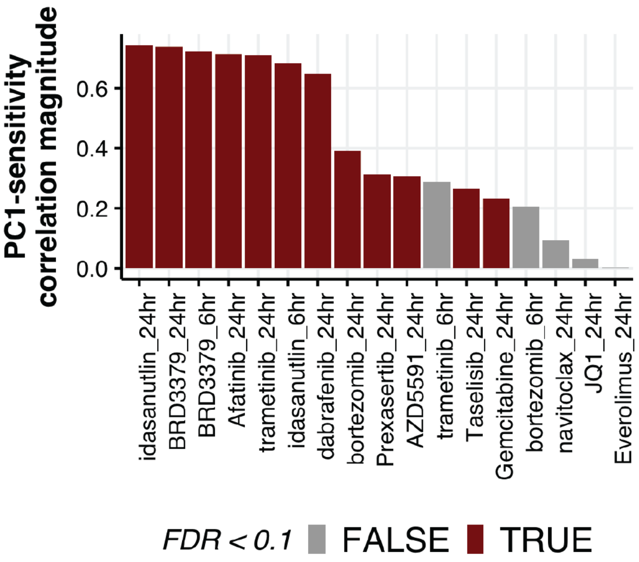


**Supplementary Fig. 13: Top principle component of transcriptional responses is well correlated with variation in drug sensitivity.**

Pearson correlation between each cell line's measured drug sensitivity and the projection of its transcriptional response profile onto the first principle component (PC1), computed for each treatment (drug and post-treatment time point). Significant correlations (FDR < 0.1) are shown in red.

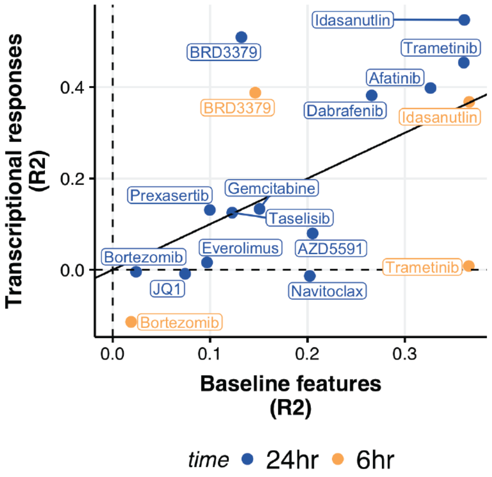


**Supplementary Fig. 14: Comparison of predictive models using transcriptional responses vs. baseline features.**

Same as **Fig. 6g**, but where the model trained on baseline features used data from all available cell lines (rather than using only the cell lines with measured transcriptional response data).

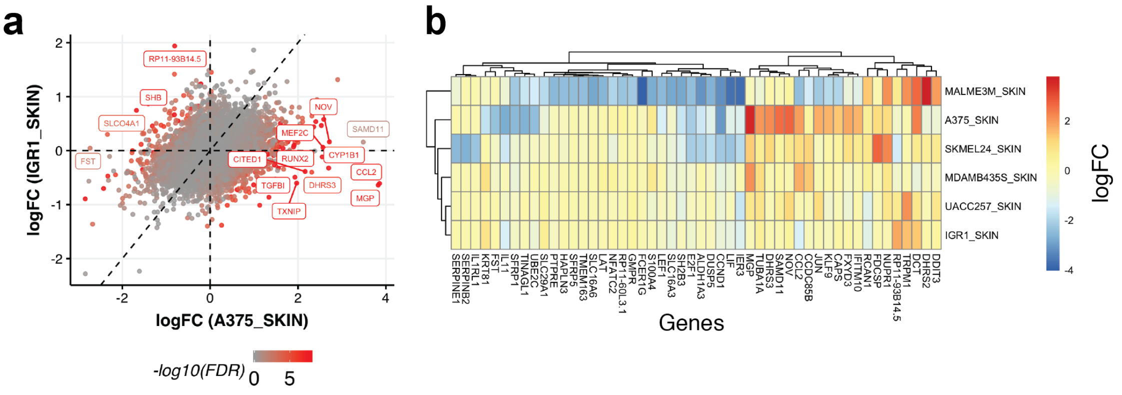


**Supplementary Fig. 15: Variability of dabrafenib responses among BRAF mutant melanomas.**

**a)** Scatterplot comparing the average response of IGR1 and A375 cells to dabrafenib treatment (24 hour post-treatment) across genes. Genes with significantly different responses between the two cell lines are indicated in red. **b)** Heatmap showing dabrafenib responses across six BRAF mutant melanoma lines for the top 50 genes with most variable responses. LFC estimates are computed with a pseudo-count parameter of 10 to stabilize estimates for low-abundance genes.

### References

Corsello, S.M., Nagari, R.T., Spangler, R.D., Rossen, J., Kocak, M., Bryan, J.G., Humeidi, R., Peck, D., Wu, X., Tang, A.A., Wang, V.M., Bender, S.A., Lemire, E., Narayan, R., Montgomery, P., Ben-David, U., Chen, Y., Rees, M.G., Lyons, N.J., McFarland, J.M., Wong, B.T., Wang, L., Dumont, N., O'Hearn, P.J., Stefan, E., Doench, J.G., Greulich, H., Meyerson, M., Vazquez, F., Subramanian, A., Roth, J.A., Bittker, J.A., Boehm, J.S., Mader, C.C., Tsherniak, A. and Golub, T.R. 2019. Non-oncology drugs are a source of previously unappreciated anti-cancer activity. *BioRxiv*.

Iorio, F., Knijnenburg, T.A., Vis, D.J., Bignell, G.R., Menden, M.P., Schubert, M., Aben, N., Gonçalves, E., Barthorpe, S., Lightfoot, H., Cokelaer, T., Greninger, P., van Dyk, E., Chang, H., de Silva, H., Heyn, H., Deng, X., Egan, R.K., Liu, Q., Mironenko, T., Mitropoulos, X., Richardson, L., Wang, J., Zhang, T., Moran, S., Sayols, S., Soleimani, M., Tamborero, D., Lopez-Bigas, N., Ross-Macdonald, P., Esteller, M., Gray, N.S., Haber, D.A., Stratton, M.R., Benes, C.H., Wessels, L.F.A., Saez-Rodriguez, J., McDermott, U. and Garnett, M.J. 2016. A landscape of pharmacogenomic interactions in cancer. *Cell* 166(3), pp. 740–754.
